# A transient neural code for feedback-driven motor corrections during reaching

**DOI:** 10.1101/2025.03.21.644350

**Authors:** Nina Kudryashova, Cole Hurwitz, Matthew G. Perich, Matthias H. Hennig

## Abstract

Movement is the result of complex, dynamic interaction between cortical and subcortical circuits. These dynamic interactions implement both feedforward motor control, arising from preparatory states, and feedback control, triggered by unexpected sensory events during movement. We show that the neural responses for feedback-driven control can be transient and small in variance, posing difficulties for unsupervised inference methods. We thus propose the Behavior-Aligned Neural Dynamics (BAND) model, which exploits semi-supervised learning to extract latent dynamics that predict both feedforward planned movement and unplanned feedback corrections. Our analysis suggests that motor corrections during movement 1) are encoded on the population level in small neural variability in primary motor (M1), but not dorsal premotor (PMd) cortex; 2) are transient; and 3) are driven by sensory feedback. Our work highlights the importance of targeted closed-loop aware methods to extract and study neural dynamics underlying complex behavioral phenomena.

---

Movement planning is a central source of variability for both neural activity and behavior during movement execution [1]. As a result, neural population activity during movement can be approximated with autonomous latent dynamics that unfold from an initial preparatory state [2]. It is hypothesized that during the preparatory phase, the motor cortex receives control inputs from the thalamus that optimize this initial state in a way that minimizes the prospective motor error [3]. The dynamics in the thalamocortical loop that unfolds from the optimized initial state orchestrates the sequence of muscle contractions that perform the feedforward control of the planned movement in the absence of any unexpected perturbations to behavioral outcomes. Accounting for any unexpected movement perturbations requires continuous feedback-driven control based on sensory input.

Sensory feedback can perturb the dynamics in the thalamocortical loop during movement execution. These perturbations can be inferred from neural activity as a solution for an optimal control problem [4, 5], provided that they significantly alter the neural population trajectory. This assumption might not hold for feedback control of movement, since the feedback-driven corrections may only cause transient, short-lived deviations from the movement plan (Fig. 1a). Such transient neural dynamics comprise a small fraction of recorded neural variability, even if it corresponds to large deviations in movement trajectory (Fig. 1b).

**Fig. 1.**
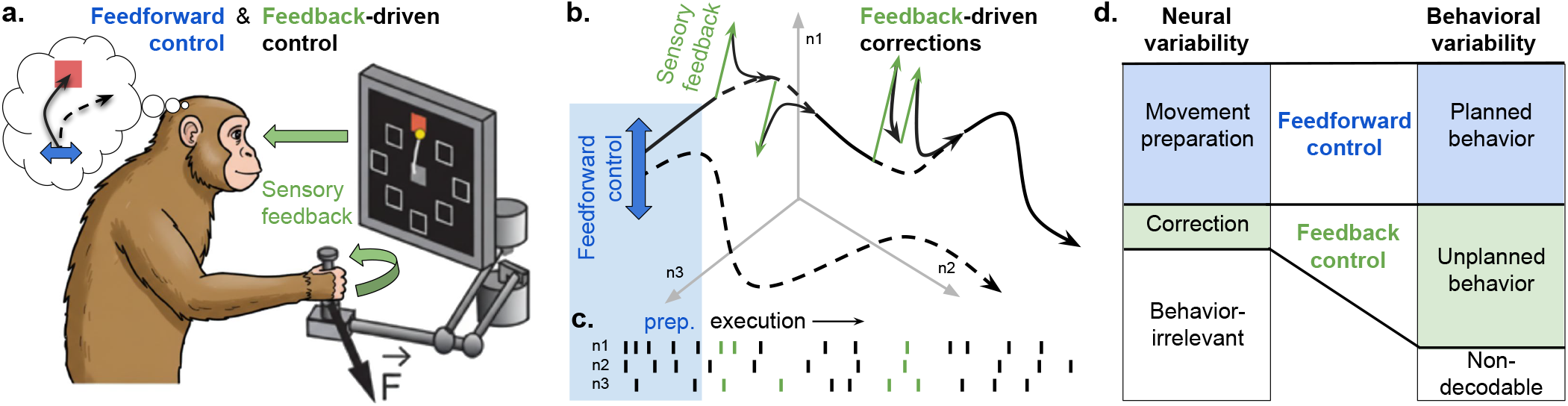
Main hypothesis: neural dynamics for feedback-driven movement corrections are transient. a) A monkey performing center-out reaches with a force field (reproduced from [6]) uses two motor-control strategies: feedforward (blue) and feedback (green). b) Neural population activity in M1 explained in a dynamical systems framework. Feedforward, or open-loop, control sets the initial condition (blue) for motor cortical dynamics; the dashed line shows a baseline trajectory, and the solid line – a trajectory after feedforward adaptation. Feedback, or closed-loop, control further adjusts the trajectory during movement execution based on sensory input (green). c) Neural responses signaling feedforward-controlled long-timescale changes in the neural population state (black) vs. feedback corrections (green). If feedback-driven responses are transient and short, they manifest in a very few spikes. d) As a result, a small fraction of neural responses (green) can execute feedback-driven control to correct large unplanned deviations in behavior.

In this paper, we demonstrate that feedback-driven movement corrections are indeed encoded in small neural variability. We analyze recordings from the primate motor cortex during a center-out reaching task with a force-field perturbation. We demonstrate that corrections during movement are encoded at the population level in primary motor (M1), but not dorsal premotor (PMd) cortex. We develop a method that discovers the neural code for feedback-driven movement corrections, and show that they result in transient deviations from the latent trajectory. We also show that this movement correction signal is usually discarded by unsupervised latent variable models as noise. Finally, the causal structure of our model allowed us to analyse temporal relationships between neural dynamics and behavior, demonstrating that sensory feedback that provides the motor error signal in perturbation trials reaches M1 with a 90 ms delay.

## Results

To test the hypothesis that feedback-driven movement corrections are encoded in small neural variability, we analyzed neural recordings from monkeys performing a center-out reaching task with a force field perturbation (Fig. 2a) [6]. In this task, monkeys were required to move a cursor on the screen using a manipulandum, performing reaches to one out of eight specified targets. In perturbed trials, a force was applied on the manipulandum perpendicular to the hand velocity of the monkey. The perturbation caused deviations from a planned trajectory, necessitating online motor correction based on tactile and visual feedback. Each experimental session consisted of three epochs: unperturbed Baseline (BL) trials, followed by Adaptation (AD) trials with the perturbation, followed by Washout (WO) trials in which perturbation was removed and the monkey had to re-adapt to normal movements. The trajectories towards the target were straight in BL trials and curved and distorted both in AD and WO trials (Fig. 2a).

**Fig. 2.**
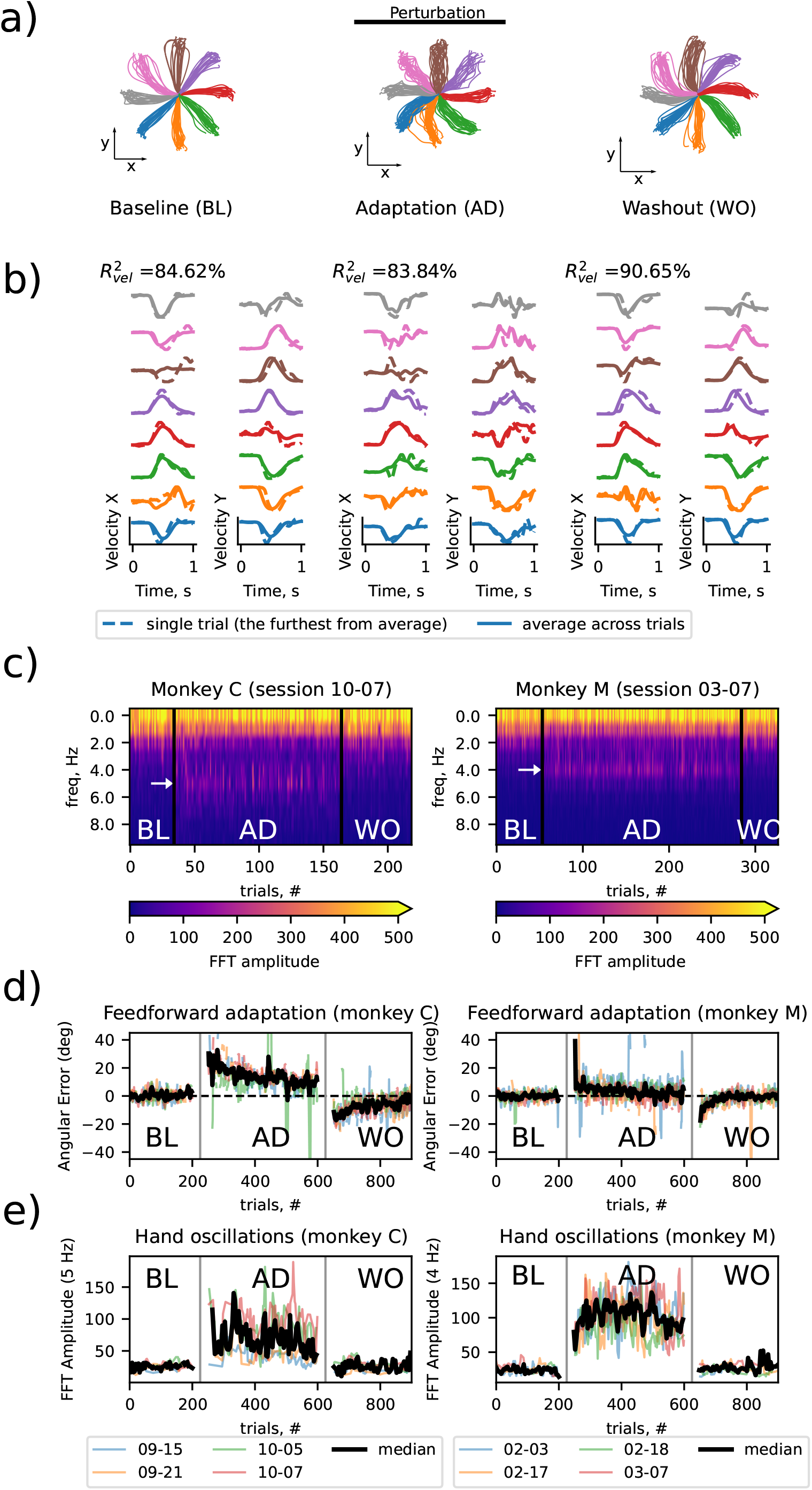
Hand velocity during center-out reaches with a force-field perturbation exhibited 4-5 Hz oscillations. **a)** Hand position during baseline (BL), adaptation (AD) and washout (WO) trials shows the effect of perturbation on hand trajectories **b)** Hand velocity in corresponding epochs; the trials that deviate the furthest from th3e trial-average velocity for a given condition are shown. **c)** Fourier spectrum across epochs in the last session for each monkey; white arrow shows the peak oscillating frequency. **d)** Angular deviation of the hand reaching trajectory at peak velocity (4-trial running average; replicates methods in [6]); **e)** FFT Amplitude (4-trial running average) of the mode corresponding to the peak oscillating frequency across epochs; sessions are color-coded.

### Feedforward motor adaptation vs. feedback control during reaching in a force field

Previous studies [6–10] have focused on the feedforward motor adaptation: adjusting the initial movement direction across trials. However, this strategy alone cannot compensate for unstable, velocity-dependent perturbations like a force field. Feedback-driven control can refine the hand trajectory, but inherent sensory and motor delays typically cause oscillations. We observed such high-amplitude hand velocity oscillations in perturbed trials (Fig. 2b, middle). The amplitude of oscillations is comparable to the overall change in velocity during reaching towards the target. These oscillations have a period of approximately 200 to 250 ms, or frequency of 4-5 Hz, depending on the animal (Fig. 2c). They appear in adaptation epochs of all recording sessions in both subjects, but vary in magnitude between sessions (Figs. B1-B2).

An analysis of the recorded behavior indicates two control strategies were used. First, we analyzed the angular deviation from the average baseline trajectories towards the target (Fig. 2d). As previously reported [6], the monkeys initially exhibited large angular errors upon introduction to the force field, which gradually declined until behavior stabilized. The errors increased again when the force field was turned off (Washout), yet with an opposite sign. Such a persistent change in feedforward control policy is the hallmark of motor adaptation. In contrast, the amplitude of the velocity oscillations showed a different time course (Fig. 2e). It remained high while the force field was on, and diminished as soon as the perturbation was removed. This result suggests that the monkeys combined two distinct control strategies, feedforward adaptation and feedback control, to adjust their reaching in a force field.

### PMd encodes overall hand trajectory while M1 additionally encodes motor corrections

The presence of strong hand velocity oscillations in some sessions provides the opportunity to find their neural correlate in motor cortical areas. We trained a decoder (a bi-directional RNN) to predict behavior from the neural activity, which takes forward and backward passes through neural data in order to account for both feedforward and feedback control of behavior. We used trial-average reaches to each target as a baseline model with no feedback control (Fig. 2b, *R*^2^=83.84%; more sessions in Table H3). The decoder, in contrast, decoded a considerable portion of additional trial-to-trial variability in hand velocity (Fig. 3a, *R*^2^ = 92%; more sessions in Table H4).

**Fig. 3.**
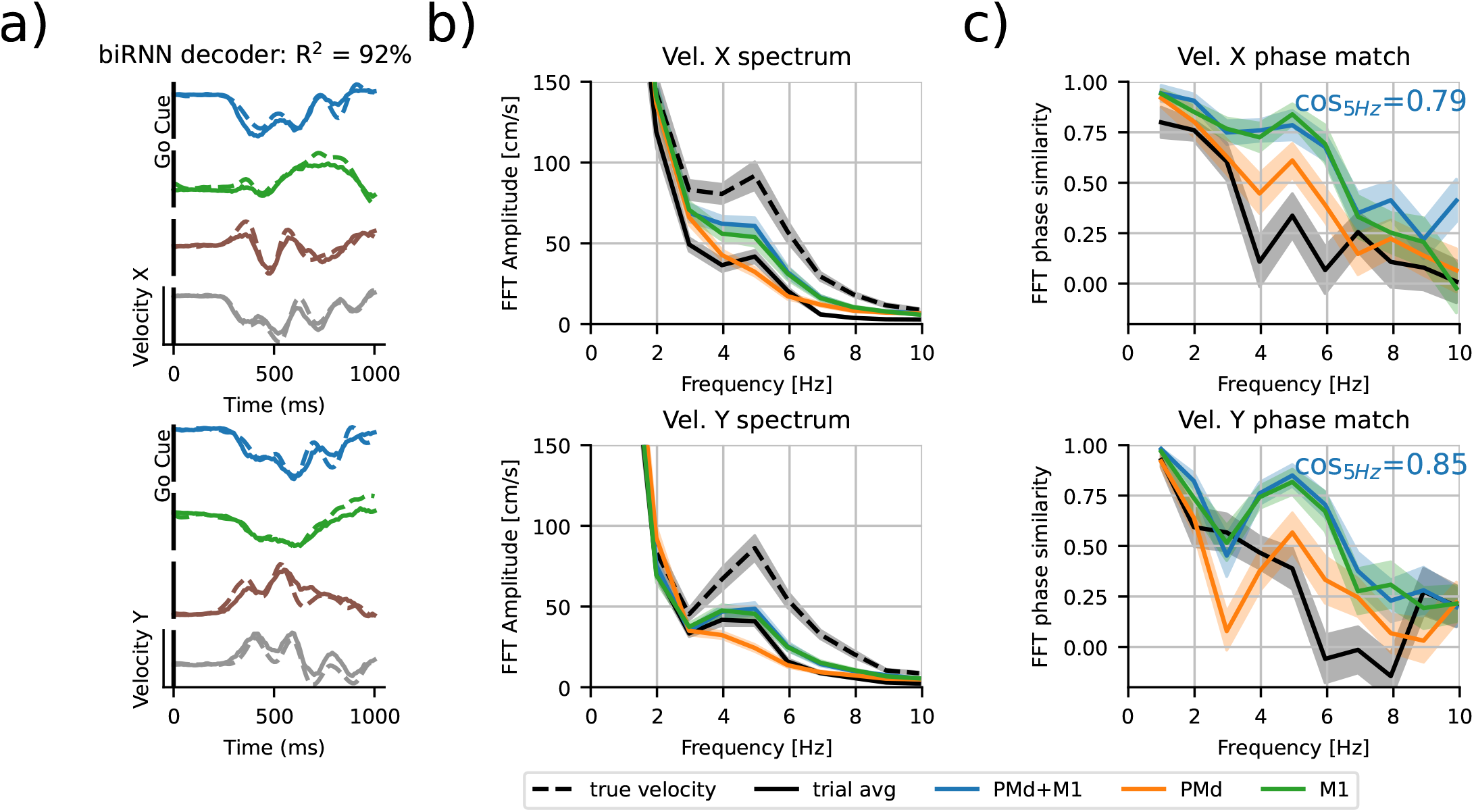
Hand velocity oscillations are decodable from neural activity in M1, but not PMd. **a)** Velocity predicted by a supervised bi-directional RNN decoder in example AD trials (solid line – model prediction from PMd+M1, dashed – ground truth) **b)** Comparison of Fourier spectra of hand velocity predictions (from both or either brain areas) and of recorded hand velocities. **c)** Cosine similarity between Fourier modes of true velocity vs decoded velocity; the similarity at 5 Hz indicates whether the phase of oscillations is correctly captured.

To locate the neural code for hand oscillations, we compared the Fourier spectra of hand velocities decoded from different brain areas. We observed that a prominent peak at 5 Hz was correctly predicted only when decoding from M1 responses, but not from PMd responses (Fig. 3b). Importantly, the phase of the oscillations, which varies between trials, was only decodable from M1, but not from PMd (cosine similarity *cos*_5*Hz*_ = 0.85 for M1, *cos*_5*Hz*_ = 0.60 for PMd; Fig. 3c; for other sessions see Fig. H13). This indicates that hand velocity oscillations are due to feedback control and triggered by sensory feedback, which is processed by M1, but not PMd [11].

While oscillations were decodable from M1, we saw very few individual M1 neurons exhibiting the corresponding oscillations (see Appendix A), suggesting that this information is encoded on a population level. The overall direction of movement was decoded from PMd and M1 almost equally well, although the velocity decoding performance was always higher for a combination of two areas (Table H4). Together these results suggest that PMd and M1 carry complementary information about movement.

### Behavior-Aligned Neural Dynamics model disentangles feedforward vs. feedback-driven latent dynamics

To capture both feedforward and feedback motor control, we introduce the Behavior-Aligned Neural Dynamics (BAND) model. BAND employs latent dynamics to model neural population activity, ensuring high-quality neural reconstruction, and, as a secondary objective, aligns latent representation to behavior [12]. Its core is a sequential autoencoder, which is based on the well-established LFADS model [5], and infers single-trial latent dynamics and control inputs from neural activity (Fig. 4). Despite the high quality of the neural reconstruction, LFADS does not guarantee joint identifiability of latent dynamics and control inputs, and, therefore, feedforward dynamics and feedback control remain entangled in the latent space. To disentangle feedforward and feedback control, BAND imposes stronger constraints on the encoder networks. An initial condition (*g*_0_) encoder can only access spiking activity in preparatory period, while a control input (*u*(*t*)) encoder accesses data from movement onset up to the current time *t*. This constraint results in a *causal* encoding model where only past activity contributes to future predictions. As a result, the initial conditions capture the feedforward movement plan, while control inputs *u*(*t*) capture potential behavior feedback.

**Fig. 4.**
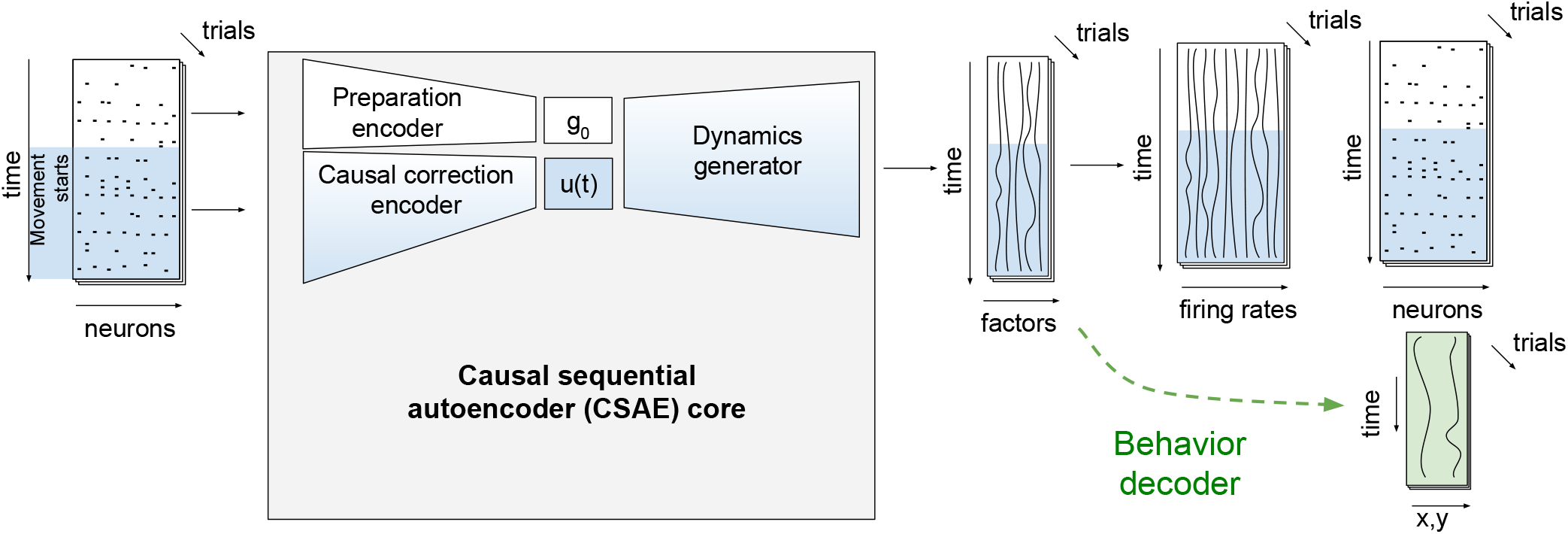
Our **B**ehavior **A**ligned **N**eural Dynamics model (BAND), built on top of a causal sequential autoencoder (CSAE), uses a behavior decoder from latent factors (green) to capture small behaviorally-relevant neural variability. The blue region, highlighting the movement execution phase, is encoded separately from preparatory activity, disentangling feedforward and feedback-driven latent dynamics components during movement execution.

The additional supervision in BAND (Fig. 4, green) ensures that behavior-related neural variability can be captured even if they cause only a small change in neural activity. We used a sequence-to-sequence linear decoder (from [12]; see Methods), trained jointly, end-to-end, with the core causal sequential autoencoder (CSAE) model. This decoding approach captures complex temporal relationships between neural activity and behavior, which we anticipated from a mixture of feedforward and feedback-driven control of movement.

The structure of our model now allows systematic hypotheses testing by manipulating its components. We first found that preparatory activity alone correctly predicts the reach direction, as the initial conditions capture the ring structure of the task with or without behavior supervision (Fig. 5a).

**Fig. 5.**
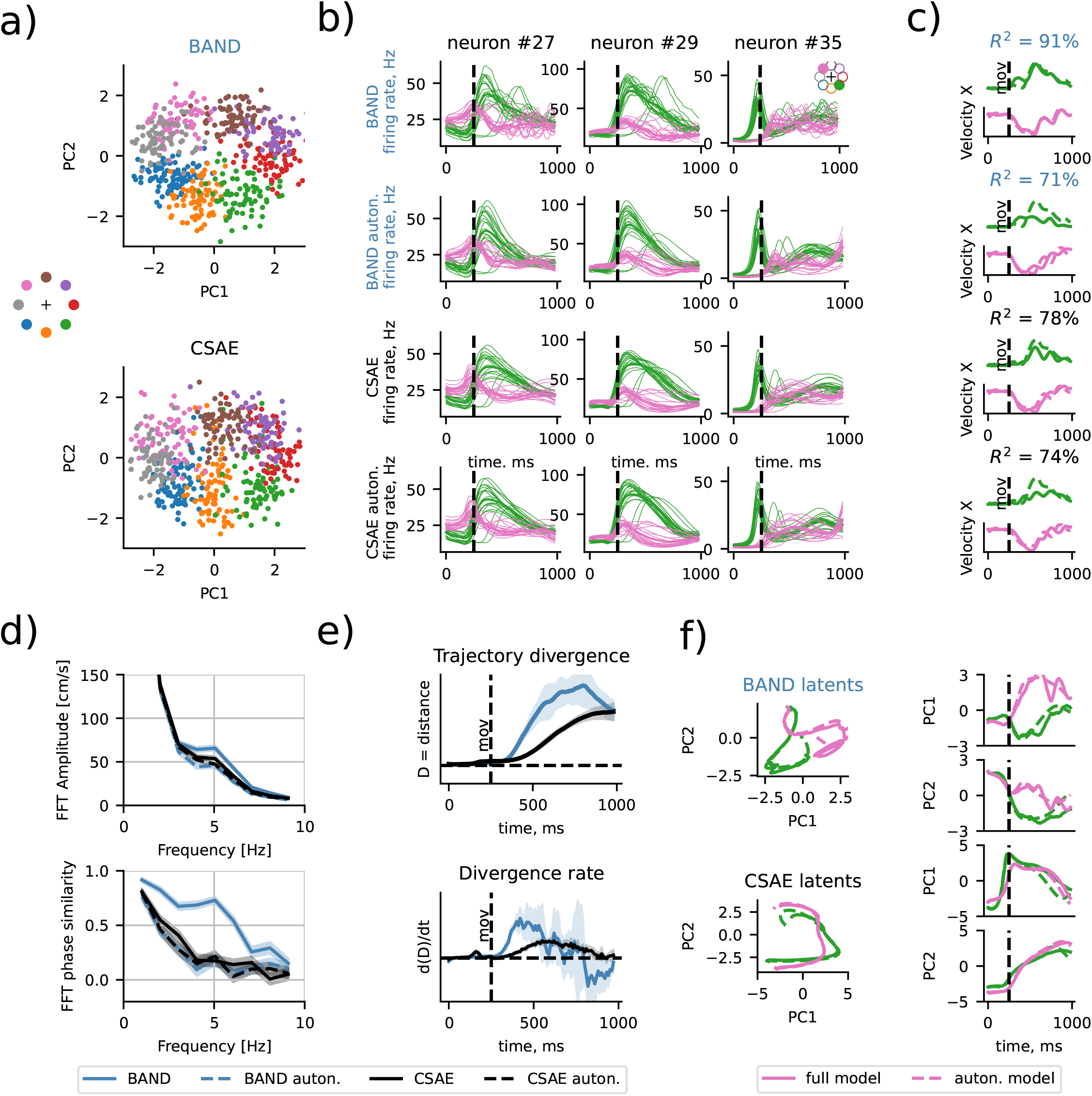
Model-based analysis confirms movement corrections due to feedback control, which makes a small contribution to neural variability while significantly improving behavior reconstruction (*R*^2^). **a)** Initial conditions capture the ring structure of the task, with (BAND) or without (CSAE) behavior supervision; **b)** Control inputs add oscillations to predicted firing rates; responses to two example targets (upper-left pink, bottom-right green) for three example M1 neurons are shown; the unsupervised baseline model (CSAE) does not capture oscillations in firing rates, despite having a controller. **c)** Behavior reconstruction in a full BAND model (top) is considerably higher (*R*^2^=91%) than that of an autonomous BAND model (71%) and unsupervised baseline with a controller (78%) or without (74%). **d)** Fourier analysis shows that only BAND with a controller captures the phase of the oscillation (cosine similarity at 5 Hz is 0.83). **e)** The BAND controller causes transient deviations in latent trajectories during movement, while CSAE trajectories diverge towards the end. Note that the magnitude of divergence in the latent space is arbitrary and does not proportionally translate into behavior prediction. **f)** Example latent trajectories for 2 trials shown in (b-c).

To understand the role of the feedback controller, we disabled this pathway in BAND after training. Then the predicted oscillations in firing rates of some M1 neurons disappeared. These predictions from feedforward dynamics in BAND were qualitatively similar to the firing rates predicted by CSAE (Fig. 5b). This suggests that behavior supervision in the full BAND model does not affect the autonomous neural dynamics (Fig. 5a), but instead introduces additional behaviorally-relevant neural variability to the controller inputs (Fig. 5b).

The quality of behavior prediction decreased significantly when the control input was removed: from *R*^2^ = 91% to *R*^2^ = 71% (Fig. 5c). In comparison, linear sequence-to-sequence regression from CSAE factors resulted in *R*^2^ = 78%, which is only slightly better than that of the autonomous BAND model. This result is unsurprising since the BAND model without controls has a lower capacity in its latent space due to ablation of control inputs and, therefore, is expected to perform worse than CSAE with a controller. The reach direction was still captured after ablation of control inputs, yet the hand velocity oscillations were out of phase, similar to behavior decoded from CSAE factors (Fig. 5d).

We next analyzed the divergence of the latent trajectories induced by the controller. We found that controlled trajectories both in BAND and CSAE diverge initially at movement onset, correcting for the speed of movement (see Fig F11). However, BAND trajectories then stop diverging and begin converging to the autonomous trajectory towards the end of the trial (Fig. 5e). During movement, BAND trajectory oscillates around the autonomous trajectory (Fig. 5f). In contrast, CSAE trajectories persistently diverge throughout the trial. We also isolated inferred point perturbations and confirmed that the resulting deviations are transient (Supplementary Fig. E9). This suggests that BAND with behavior supervision is indeed capable of capturing transient behaviorally-relevant changes in latent dynamics.

### Movement corrections are encoded in a small portion of neuronal variability

Capturing movement correction information in our BAND model did not improve the neural reconstruction in comparison to the unsupervised CSAE model (Fig. 6a), with both models accounting for 0.22 bits/spike. Ablating the controller in BAND did not significantly change the neural reconstruction quality (Fig. 6a), resulting in only 0.03 bits / second change. We estimated the lower bound on the total trial-to-trial variability, using BAND as the best estimator of total neural variability. We found that additional variability captured by the BAND controller, which corresponds to feedback-driven corrections, comprises at most 31% of total trial-to-trial variability.

**Fig. 6.**
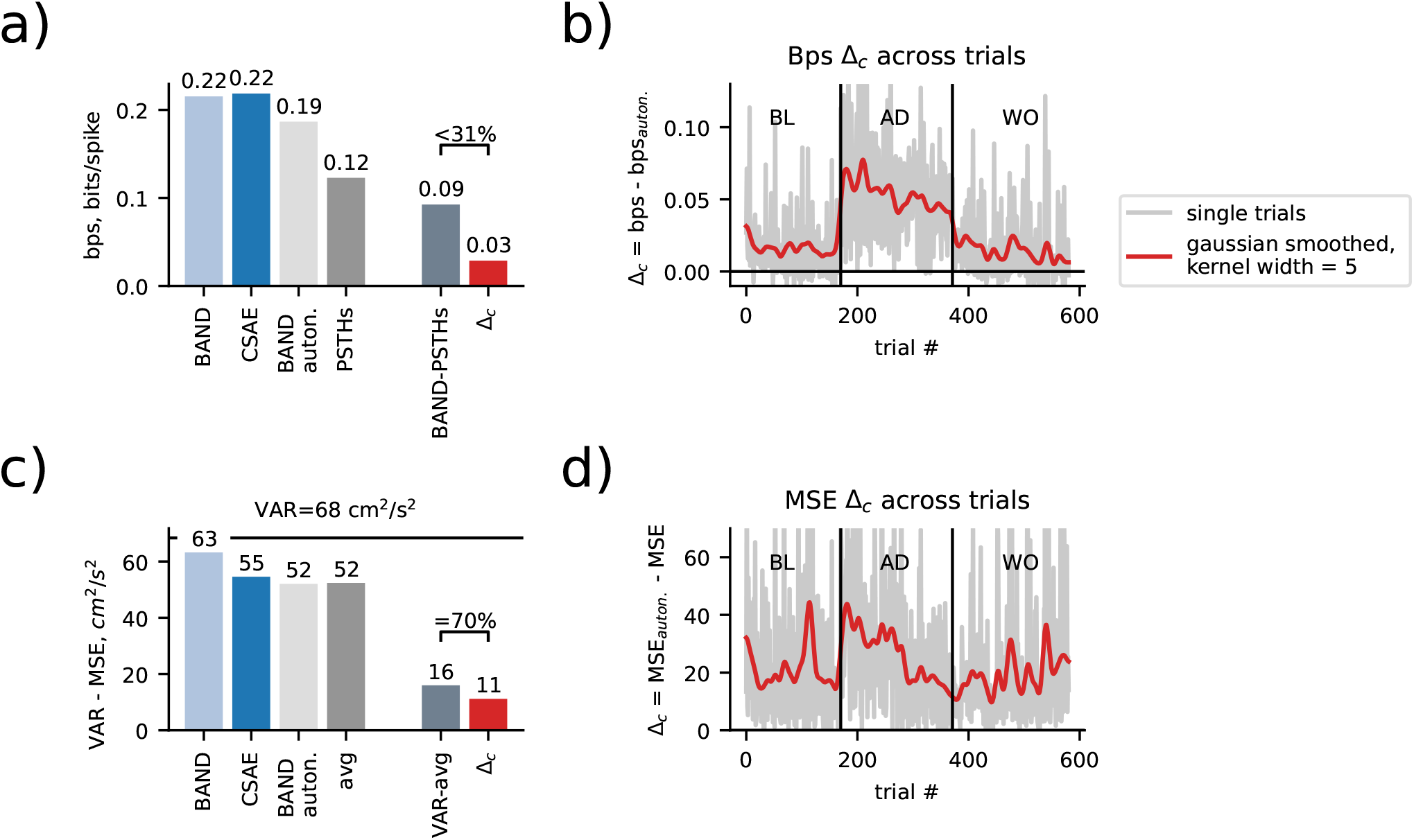
Control inputs account for small neural, yet significant behavioral variability. **a)** A comparison of neural reconstruction quality between the full BAND model, unsupervised baseline model (CSAE), an autonomous BAND model (with ablated controls) and the neural reconstruction based on the average firing rate for every reach direction. The difference between full and autonomous BAND models (Δ_*c*_, red) comprises at most 31% of trial-to-trial variability in neural activity. **b)** Neural variability explained by BAND controller across different trials and epochs. **c)** Same as (a), but comparison of behavioral reconstruction instead of neural reconstruction. Explained variance (total variance minus mean squared error) is used as a measure of reconstruction quality. BAND controller accounts for 70% of all behavioral trial-to-trial variability. **d)** Behavioral variability explained by BAND controller across different trials and epochs.

Note that the autonomous BAND model could still predict much more neural variability compared to the variability captured with condition-average peristimulus-time histograms (avg PSTHs 0.12 bits/spike, Fig. 6a, left), suggesting that initial conditions capture some trial-to-trial variability (e.g. feedforward motor adaptation). We found that neural reconstruction explained by input-driven dynamics remained high throughout the adaptation epoch, suggesting that controller input accounted for feedback-driven responses to force-field perturbation (Fig. 6b). Indeed, this additional neural variability modelled with the controller follows a similar across-trial trend as the power of 5-Hz neural oscillations in hand velocity (see Supl. Fig. F10). This result confirms that preparatory activity is a major source of neural variability [1] (avg PSTHs), while feedback-driven corrections are encoded in patterns of neural activity with small variability (Δ_*c*_).

Likewise, preparatory activity was confirmed to be the main source of behavioral variability, accounting for more than 75% of total behavioral variability: the autonomous BAND model accounted for 52 out of 68 *cm*^2^*/s*^2^ (Fig. 6c). The condition average behavior accounted for the same amount of behavioral variability as autonomous BAND, suggesting that autonomous dynamics could account for all the target-related variability. Yet, there was still a significant fraction of decodable trial-to-trial behavioral variability, 70% of which was captured by control inputs in BAND (Fig. 6c). This additional behavioral variability followed a similar trend across adaptation trials as additional neural variability, yet there were some unperturbed trials (BL/WO) that had a strong behaviorally-relevant input from the controller. We found that these are the trials in which the movement was slower (see Fig. F11).

In summary, there is a second major source comprising 70% of trial-to-trial behavior variability (Fig. 6c). This extra variability is accounted for by changes in activity that are small compared to ongoing neural dynamics and comprise at most 31% of trial-to-trial variability in spiking activity (Fig. 6a).

### Behavioral variability captured with behavior supervision feeds back into neural activity

To analyze the relationship between neural activity and behavior, we visualized the weights of the BAND behavior decoder. Causal constraints on the inference network in BAND enable this analysis, ensuring that latent factors of the model are based only on the past spiking activity. At the same time, the acausal sequence-to-sequence behavior decoder can use the whole time course, including future activity, to predict hand velocity at each time step.

For each latent factor and each component of hand velocity, the decoder weights can be represented as a matrix (Fig. 7a). The upper triangle of this matrix corresponds to the neural factors that causally control behavior, while the lower triangle represents behavioral feedback. We visualize the weight matrix after training the model on the data aligned to movement onset time. We note two important features, typical for the all decoder weights (see full model visualization in Fig. G12a). First, there is a concentration of strong positive weights (blue) that connect the initial values of latent factor in the preparatory state (before movement begins at 250 ms) and behavior at about 250 ms after movement onset (500 ms from the start of the trial). These weights correspond to feedforward motor planning. Second, there are negative (red) weights on the lower diagonal, which connect hand velocity changes that are slightly lagging behind the neural latent factors. These weights correspond to behavioral feedback.

**Fig. 7.**
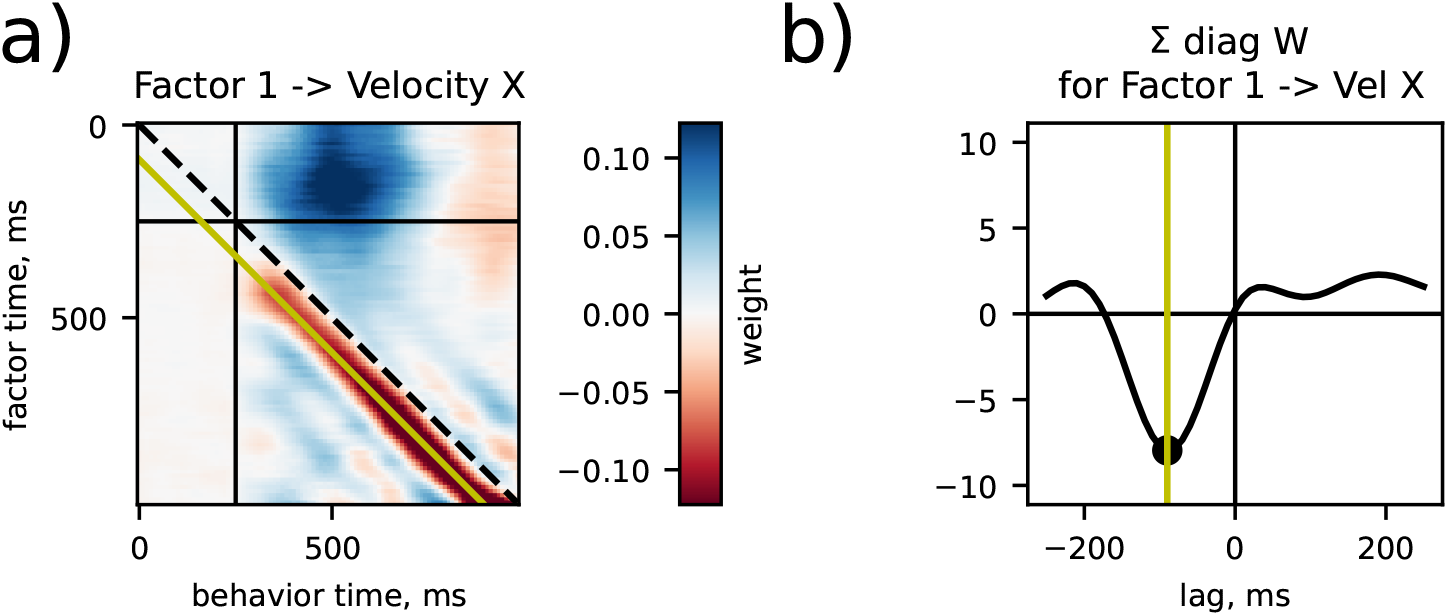
Latent factors in BAND model capture behavioral feedback with a 90 ms lag. **a)** Behavior decoder weights, transforming a latent factor (here, number 1) into hand velocity (here, component X; see all factor-behavior pairs in Fig. G12). Black vertical/horizontal lines separate preparation and movement phases. Yellow diagonal indicates 90 ms lag between behavior and the latent factor. **b)** Sum of behavioral weights in (a) across the diagonals in movement phase.

We then sum the weights along the diagonals of the weight matrices during the movement phase (lower right block of the weight matrix, shown in Fig. 7a). Each diagonal corresponds to a fixed lag between neural factors and hand velocity. The peak in these weights indicate that changes in hand velocity lag behind neural factors by 90 ms (Fig. 7b; all components are shown in Fig. G12a). This result is in agreement with the typical times required for processing sensory input [13] and is slightly slower than 50 ms feedback reported earlier for the reaction to mechanical perturbation in arm movement in monkeys [14].

## Discussion

In this work, we found that feedback-driven motor control accounts for a considerable portion of behavior variability, while being encoded in transient changes in neural activity. These transient changes comprise a small fraction of neural variability, compared to the ongoing neural dynamics. As a result, the neural code for feedback-driven motor control can be missed by unsupervised methods, which identify the most prominent patterns of neural activity without access to behavioral output. We proposed a methodology that captures both feedforward movement planning and feedback-driven correction in a disentangled representation within a single latent dynamics model.

Our results offer a revised perspective on the dynamical system view of the motor cortical neural activity. The classic view focused on feedforward control, identifying preparatory activity as the major source of both neural and behavioral variability. However, those results were obtained in highly controlled experimental settings, while natural behavior in a changing environment is likely more feedback-driven. The main paradox that we revealed is that even significant feedback-driven behavior changes correspond to small changes in neural activity during movement. These results are in line with theoretical works that view the primary motor cortex (M1) as a feedback controller [15–17], which aims to minimize prospective motor error [3] and is shaped by sensory expectation [18]. Here we also highlight a new practical implication: the neural code for feedback-driven motor control cannot be revealed by only focusing on large neural variability, and instead requires a specialized data-driven modeling strategy.

Motor cortical activity, including feedback-driven components, still has a relatively low-dimensional structure. Our BAND model required few factors (8 factors in Fig. 7) to capture dominant features of the hand reaching in force-field: target direction and feedback-driven oscillations. This confirms that the motor cortex operates on a low-dimensional manifold [19], identified not just by maximizing explained neural variability but also by maximizing explained behavioral variability.

Finally, we developed a strategy for capturing both the feedforward movement planning and feedback-driven corrections in a latent variable models. We demonstrated that this requires two modeling assumptions: first, it requires dynamics to account for the movement plan and second, it requires behavior supervision to ensure that the neural code for ongoing movement corrections is captured. We also utilized a sequence-to-sequence decoder for behavior supervision and imposed additional causality constraints on the model that disentangle feedforward and feedback-driven control withing the latent representation. As a result, specific modifications of the model allowed us to explore the relationships between neural and behavioral variability based on recorded data, identifying feedforward motor control (motor planning) and lagged feedback-driven motor control.

Capturing behavior feedback in neural activity makes BAND unique among other semi-supervised models, such as PSID or CEBRA (see Appendix C.1). Strong inductive biases in BAND distinguish it from CEBRA, making BAND components interpretable and focused on behaviorally relevant timescales. PSID belongs to the same class of dynamical system models, yet, unlike BAND, the PSID behavior decoder is restricted to predicting behavior that is temporally aligned to latent factors (with a fixed lag between the two) [20]. A more recent version of this method, dissociative prioritized analysis of dynamics (DPAD [21]), introduced nonlinear dynamics and nonlinear relationships between latent dynamics and behavior, yet retained temporal alignment between time-series as in PSID. While this alignment makes PSID/DPAD easily applicable to timeseries acquired in real time, this limitation prevents these models from capturing bi-directional temporal relationships between neural activity and behavior, which were instrumental for this study (Fig. 7). In contrast, our previous work, Targeted Neural Dynamical Modeling [12], used the same approach to sequence-to-sequence decoding as BAND, yet did not have the controller to capture feedback-driven movement corrections.

Our analysis was limited to correctly executed reaches in highly trained animals. As a result, we did not consider trial-wise error processing and motor learning that is facilitated by rewards at the end of the trial. Instead, we focused on online, continual error correction. A study by Feulner et al. (2021) [22] considered rapid adaptation to a visuomotor rotation, in which targets on the screen were shifted by 30 degrees. Using modular RNNs, they simulated neural activity changes based on the plasticity of specific modules during adaptation. Their findings suggested that motor adaptation in this context was mediated by upstream areas providing input to motor cortices. A more recent paper from Feulner et al. (2025) [23] linked online corrections and progressive adaptation to changing conditions under a single optimal control framework, yet only for the visuomotor rotation. However, the mechanisms of adaptation to visuomotor and force-field perturbations are likely different [6], since force-field perturbations directly affect the physical forces experienced by the limb, while visuomotor perturbations alter the visual feedback about the movement without changing the actual forces. Another study by Perich et al. (2024) [24] examined the role of feedback in motor learning using a naturalistic task in which monkeys reached for, grasped, and pulled objects mounted on a robotic arm. They found that motor cortex activity was strikingly input-driven surrounding behavioral error correction, and that these input-driven dynamics were isolated in a subspace of the population activity that captured somatosensory feedback. These findings were also causally validated with electrical stimulation of ascending somatosensory tracts. Further analysis with modular biological RNN models and kinematic hand models, as well as causal experimental validation, would be required to localize plasticity in response to force-field perturbation, which is beyond the scope of this paper.

In more naturalistic, unstructured experimental settings [25] we would expect a larger contribution from the feedback-driven rather than feedforward control in response to unpredictable changes in the environment. Conversely, we would expect an even bigger gap in performance between unsupervised models and semi-supervised models (such as in MC RTT in Appendix C). The core of the model (RNN generator) can be swapped for more performant architectures [26–29] to process larger and richer naturalistic data. Yet, retaining BAND’s causal constraints and behavior supervision are two principles that should guide model design to capture transient feedback control signals.

Our results demonstrated that movement corrections can be captured based only on the past neural activity, as BAND is fully causal at the inference time. Therefore, our modeling approach could also be extended (similar to [30]) to real-time applications in brain-computer interfaces (BCIs). Future work can extend BAND to identify shared latent dynamics in larger datasets with multiple animals, offering an exciting avenue for studying generalizable cross-animal neural representations [28, 31–34]. While feedforward motor control exhibits a shared pattern of activity across individuals [31], the same likely applies to the neural code for feedback control [35]. Therefore, identifying conserved patterns of neural activity with BAND can enable designing BCIs for paralyzed patients, for whom training a behavior decoder is no longer possible.

## Methods

### Neural and behavior reconstruction metrics

The quality of neural reconstruction is measured using bits-per-second (bps) metric [36]:

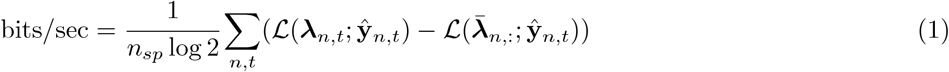

where *n*_*sp*_ is the total number of spikes,

**ŷ**_*n,t*_ is the observed spiking activity of neuron *n* at time *t*,

***λ***_*n,t*_ is the rate prediction for neuron *n* at time *t*,

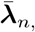,: is the mean firing rate of neuron *n*, and

ℒ (***λ***; ***y***) is the Poisson log-likelihood of observed spikes ***y*** given rates ***λ***.

Note that the CSAE/BAND models are trained to explicitly minimize ℒ (***λ***; ***y***). The idea behind this metric is to measure how much information about neural firing rates can be explained by the inferred dynamics, discounting trivial prediction of the average firing rate.

The behavioral reconstruction was measured using classic variance explained *R*^2^:

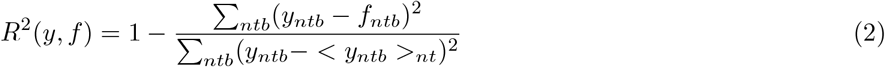

Here *y* is a behavioral variable (hand velocity for hand reaching tasks) and *f* is a model prediction; *< y >*_*nt*_ indicates an average of *y* over trial *n* and time *t* dimensions; *b* and *d* correspond to the behavioral component and target direction indices, respectively.

Note, that the *R*^2^ is often computed for each behavioral component separately and then the average *<* . *>*_*b*_ is taken (perhaps, due to the fact that such averaging over features is standard for sklearn library). This ignores the fact that components of the velocity belong to the same 2D Euclidean space. The difference between the resulting scores between our isotropic 2D *R*^2^ and *<* . *>*_*b*_ variant was negligible for the data and models presented in this paper.

In the center-out reach task, *R*^2^ is dominated by task instruction and is insensitive to the uninstructed variability. Indeed, even the simplest baseline model that uses trial average behavioral trajectory for any given direction 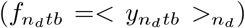 scores *R*^2^ = 84% even on the adaptation trials, and *R*^2^ = 91% on unperturbed trials (see Fig. 2b).

### Datasets with a force field perturbation

We have used 8 sessions with force field perturbation recorded from 2 Rhesus macaque monkeys (Monkey C and Monkey M) from Perich et. al. 2018 [6]. Each session consists of three epochs: baseline (BL), adaptation (AD), and washout (WO).

All sessions were used in our analysis. A few trials were excluded where velocity readings were corrupted and included unrealistically high (*>*1 m/s) numbers: 3 trials in total, 1 per session in sessions 2016-09-15, 2016-09-21, 2014-02-03.

We used 80% / 20% split between train and test sets to train and evaluate all decoders and latent variable models. We used a single random 80 % / 20% split to benchmark models (Fig. C5), and a 5-fold cross-validation to analyze BAND-model predictions across all trials (Fig. 5-7).

### Fourier analysis of hand velocity and neural oscillations

We applied Fast Fourier Transform (FFT) to the hand velocity to quantify 4-5 Hz oscillations observed empirically in adaptation trials in Fig. 2b. We applied FFT to every component of the velocity in every trial, obtaining powers (squared amplitude) of different Fourier harmonics. We then calculated the mean power across trials for each Fourier mode (shown in Fig. 2c).

We then applied a similar procedure to the firing rates of the single neurons in PMd/M1. The firing rates were estimated using a Gaussian filter with *σ* = 30 *ms*. FFT was applied to every trial separately, and then the trial average amplitude of each FFT mode was calculated. The neurons were considered oscillatory (see Appendix A) if the power of 4 Hz or 5 Hz FFT modes (the ones closest to 4 Hz and 5 Hz) in adaptation but not baseline trials was higher than mean + 4SD of the power distribution obtained from shuffled data. For each trial, spike counts were independently shuffled 500 times. The firing rate and FFT were then recomputed for each shuffled dataset. This criterion specifically identified neurons exhibiting 4-5 Hz oscillations *induced* by the perturbation, excluding neurons oscillating at these frequencies regardless of trial type. In other words, the amplitudes of the 4-5 Hz modes were required to fall within the chance distribution during baseline trials but significantly outside this distribution during adaptation trials.

### Angular deviation of hand reaching trajectory

Angular deviation in Fig. 2d was measured between hand velocities at the moment when the speed (norm of the velocity) is the highest (approximately 300 ms after movement onset). We subtracted the average angular deviation for each target in Baseline trials, to account for the natural curving of the unperturbed trajectories.

### Models of motor cortical activity and behavior decoders

#### Average hand trajectory per target

The average hand trajectory per target was calculated on the training set, containing 80% of trials. The variance explained by this average hand trajectory was evaluated with the *R*^2^ (see eq.2) on the remaining 20% trials (Table H3). These values quantify how much variability in hand velocity is explained by the difference in the reach target.

#### Bidirectional RNN decoder

We used a bidirectional RNN decoder in order to test whether hand velocity oscillations in response to perturbations are decodable from neural populations (M1, PMd, or both areas together). The architecture and hyperparameter values can be found in Table 1.

**Table 1.** Summary of the biRNN Decoder Network Architecture.

| Layer | Size / Operation |
| --- | --- |
| Input Dropout | Dropout with rate 0.3 |
| Bidirectional LSTM | Input size: number of neurons<br>Hidden size: 128<br>Number of layers: 1<br>Output size: $2 \times 128$ |
| ReLU Activation | - |
| Batch Normalization | - |
| Fully Connected (FC) Layer 1 | Input size: 256<br>Output size: 128 |
| ReLU Activation | - |
| Batch Normalization | - |
| Dropout | Dropout with rate 0.2 |
| Fully Connected (FC) Layer 2 | Input size: 128<br>Output size: number of behavior components (2) |

#### CSAE as a baseline latent variable model

We used a PyTorch implementation of LFADS model [5] to build the causal sequential autoencoder (CSAE) (code will be made available on GitHub upon acceptance). Specific changes with respect to standard LFADS include:

1. Encoding initial conditions only from the spiking in preparatory period;
2. Encoding control inputs *u*(*t*) only from the spiking in movement period;
3. Restricting the controller and its encoder to be **causal**: *u*_*t*_ depends only on spiking from movement onset to current time *t*.

These additional constraints on the inference network enabled analysis of feedforward motor adaptation vs. feedback control.

The negative loss of the model is defined as follows:

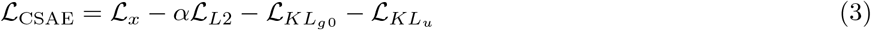

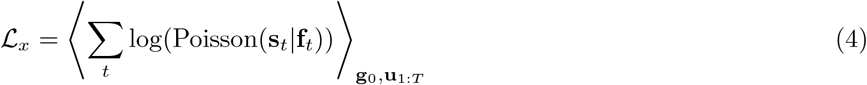

where **s** are observed spikes, **f** is a predicted firing rate; **g**_0_ and **u**_1:*T*_ are latent variables: dynamics initial condition and inputs, respectively.

We used CSAE models with 100 latent factors, 4 control inputs, 200 generator factors, and 64 encoder dimensions for both initial conditions and controller encoders. The hyperparameters were optimized using PBT training from AutoLFADS [37].

#### Sequence-to-sequence Ridge regression (SSRR) decoder

After CSAE model was trained, we trained a sequence-to-sequence Ridge regression decoder to decode hand velocity. The latent factors (dimensions: `[trials, time, factors]` ) were flattened as `[trials, time x factors]` . As a result, every trial constitutes a sample, and the whole latent trajectory provides features for the decoder.

#### BAND: Behavior-aligned neural dynamics

We propose a semi-supervised latent dynamics model called Behavior-Aligned Neural Dynamics (BAND). The main idea of the model is to use a combination of neural decoder and behavior decoder to ensure that the latent space retains the most behaviorally-relevant neural variability (Fig. 4). The whole BAND model – the latent dynamics core and the decoder – was trained end-to-end.

This idea is related to previously proposed PSID [20] and TNDM [12] methods, in which latent space was split into behaviorally-relevant and behaviorally-irrelevant parts. Yet, BAND does not make such separation, since it aims to retain rather than isolate behaviorally-relevant variability.

We constrained the architecture of the core model to enhance its interpretability, and incorporated a more expressive sequence-to-sequence decoder (Fig. 8). This decoder has the same dimensionality as the SSRR, but it is trained together with the dynamics core end-to-end.

**Fig. 8.**
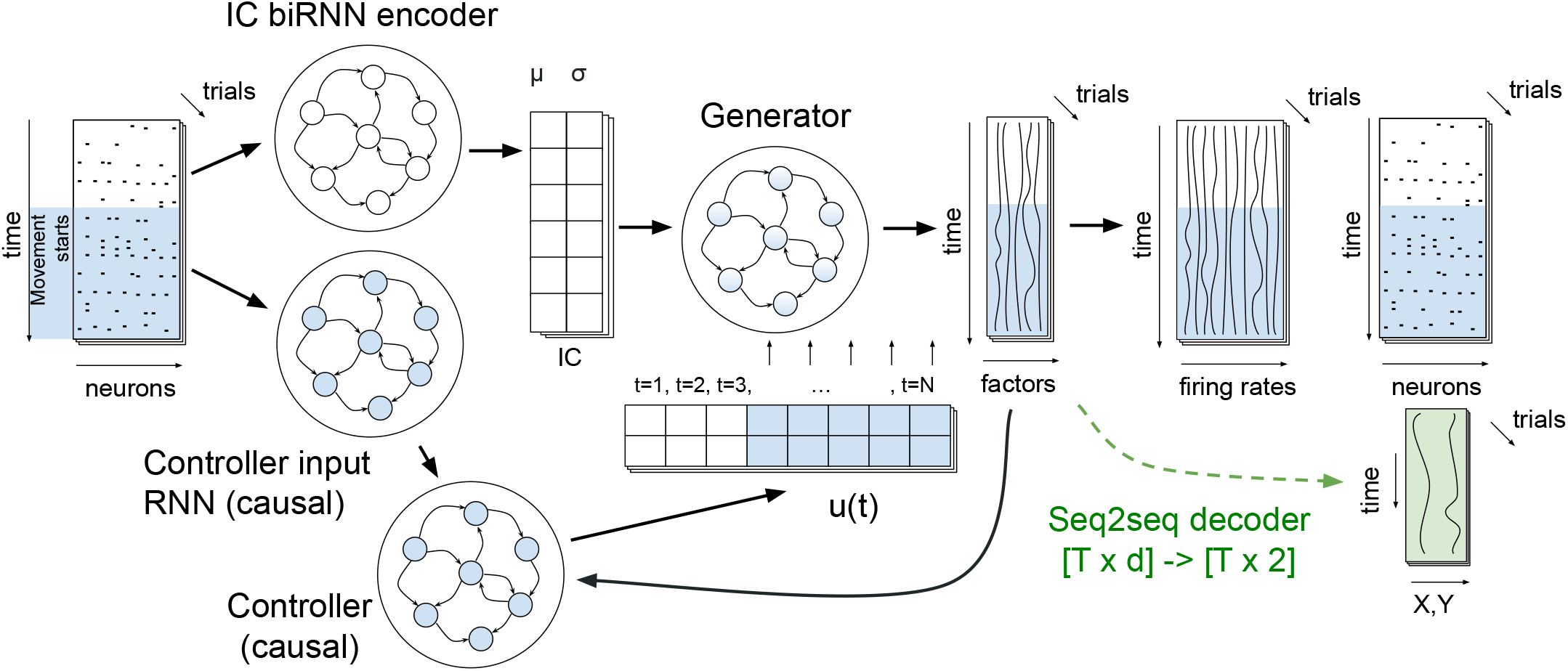
Causal BAND model architecture with enhanced interpretability. The latent dynamics inference is causal during movement (highlighted in blue).

It is important to note that while encoders during movement and dynamics generator are **causal**, the sequence-to-sequence behavior decoder remains **acausal**. This acausal decoder allows us to retrospectively relate past behavior to current neural activity (behavior feedback) when computing the loss. Yet, the inference of the latent factors during movement (blue in Fig. 8) at test time remains causal.

The negative loss of the BAND model is defined as follows:

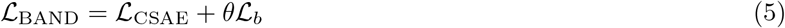

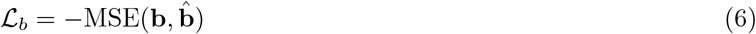

where ℒ_CSAE_ is the causal sequential autoencoder loss (eq. 3); **b** and 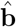 are the observed and predicted behaviors, respectively; *θ* is the weight of the behavior supervision term.

The exact value of *θ* sets the balance between prioritizing neural vs. behavioral reconstruction. If *θ* is set to a large value, it can cause the model to split the latent representation into mostly independent behavioural and neural subspaces. If *θ* is set to a small value, then the results are no different from unsupervised CSAE, which is equivalent to *θ* = 0. The intermediate values, fine tuned with hyperparameter optimization (as described below), result in high neural reconstruction and high behavioural reconstruction quality, with latent factors representing both modalities.

#### Hyperparameter optimization

We used population-based training (PBT, from AutoLFADS [37]) for optimizing hyperparameters. We used a normalized neural log likelihood (4) and behavioral *R*^2^ for all the further analysis in the paper:

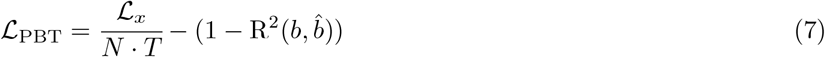

where *N* is the number of neurons and *T* is the number of time bins in a trial; ℒ_*x*_ is computed at in eq. (4), and *R*^2^ as in eq. (2).

### Trajectory divergence metric

We evaluated the divergence caused by control inputs (Fig. 5e) by measuring the distance from the perturbed state to the *closest* state on the autonomous trajectory. This metric quantifies deviations *off* the planned trajectory while discounting control inputs that only affect timing (e.g. delaying movement onset, see Fig. F11).

## Acknowledgements

The authors would like to thank Juan Gallego, Lee Miller and Robyn Greene for valuable discussions. This research was carried out with the support of the Royal Society University Research Fellowship awarded to N.K. (URF \R1 \241060), BBSRC grant (BB/X01861X/1) awarded to M.H.H., grant “chercheurs-boursiers en intelligence artificielle” from the Fonds de recherche du Quebec Santé awarded to M.P. The work of C.H. was supported in part by NIH 5U19NS107613, Simons Foundation (543023), and NSF (DBI-2229929).

## Appendix A Single neuron encoding of movement corrections

We used the Fourier transform and searched for a 4-5 Hz oscillations in neural firing rates of individual PMd and M1 neurons. We found a very small fraction of neurons with significant 4-5 Hz component in M1 (4 *±* 1%), and very few such neurons in PMd (0.2 *±* 0.5%, see Table A1). Most of the sessions did not contain any recorded oscillating PMd neurons. This suggests that M1 neurons have neural dynamic modes at a relevant timescale, and therefore the phase of oscillations in these individual M1 neurons might encode the phase of hand velocity oscillation.

Since the fraction of such neurons is small, this provides little evidence for single neurons capturing the observed oscillations. As we show in the paper based on the decoding analysis, the information about the phase and intensity of hand velocity oscillations is present in the M1 population code.

**Table A1.**
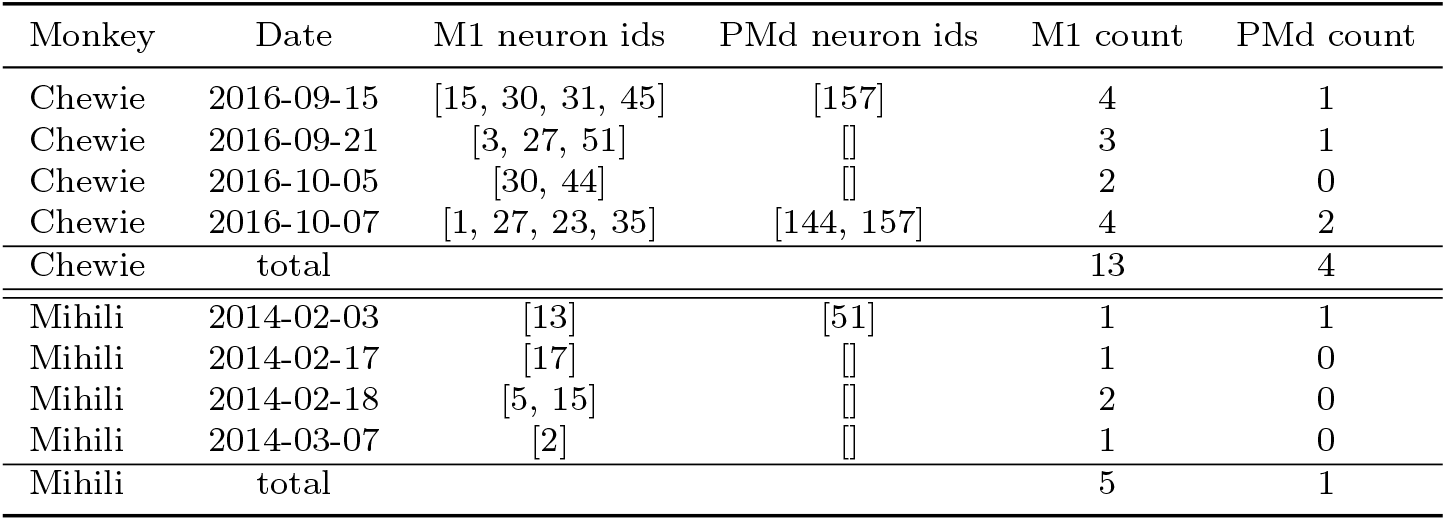
Oscillating neurons in M1 vs PMd.

| Monkey | Date | M1 neuron ids | PMd neuron ids | M1 count | PMd count |
| --- | --- | --- | --- | --- | --- |
| Chewie | 2016-09-15 | [15, 30, 31, 45] | [157] | 4 | 1 |
| Chewie | 2016-09-21 | [3, 27, 51] | ∅ | 3 | 1 |
| Chewie | 2016-10-05 | [30, 44] | ∅ | 2 | 0 |
| Chewie | 2016-10-07 | [1, 27, 23, 35] | [144, 157] | 4 | 2 |
| Chewie | total |  |  | 13 | 4 |
| Mihili | 2014-02-03 | [13] | [51] | 1 | 1 |
| Mihili | 2014-02-17 | [17] | ∅ | 1 | 0 |
| Mihili | 2014-02-18 | [5, 15] | ∅ | 2 | 0 |
| Mihili | 2014-03-07 | [2] | ∅ | 1 | 0 |
| Mihili | total |  |  | 5 | 1 |

## APPENDIX B Hand velocity oscillations can be found in adaptation trials of all sessions

**Fig. B1.**
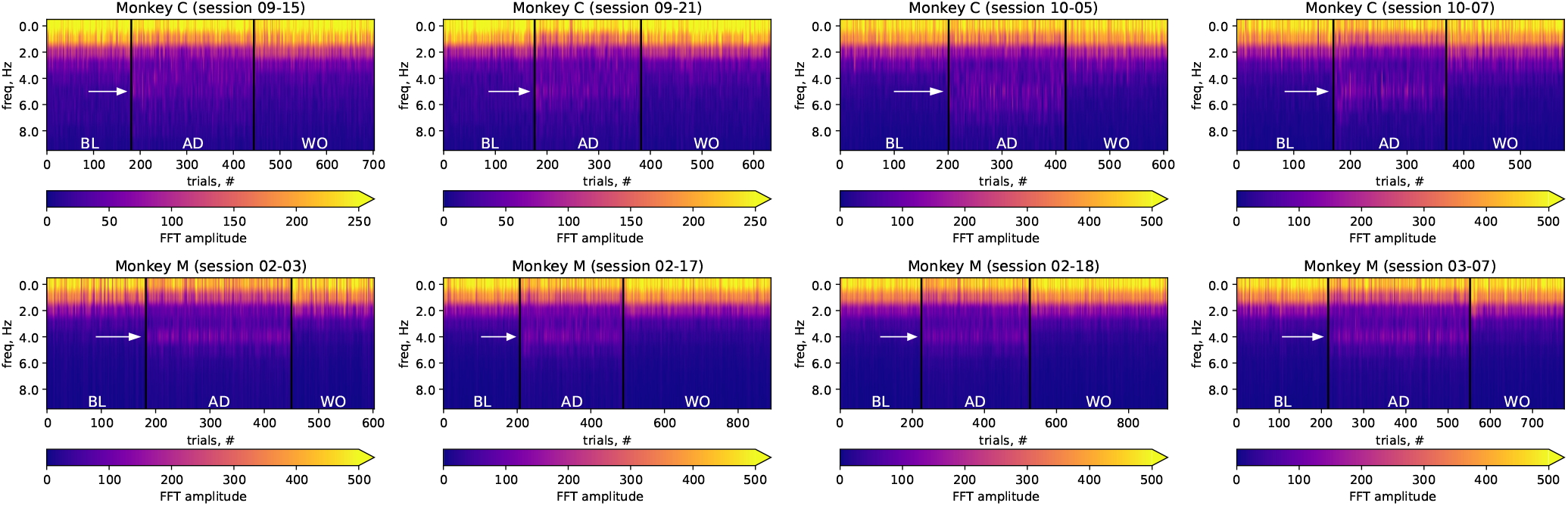
Fourier spectra of hand velocity across trials and epochs, for both monkeys and all recording sessions. The white arrow points at the oscillatory frequency (5 Hz for Monkey C, 4 Hz for Monkey M), which appears in adaptation trials.

**Fig. B2.**
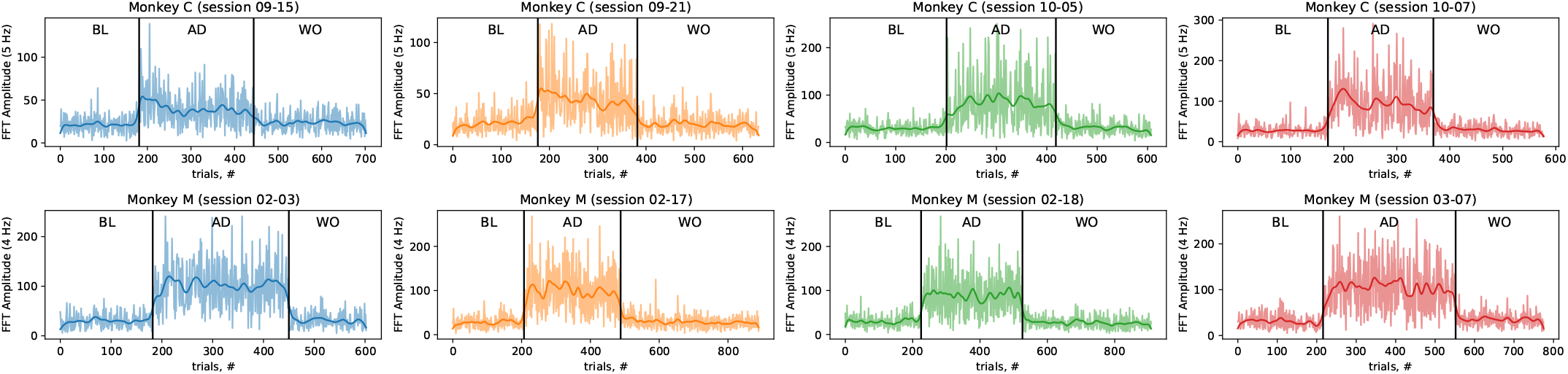
Fourier amplitude of hand velocity oscillation across trials and epochs, for both monkeys and all recording sessions.

**Table B2.**
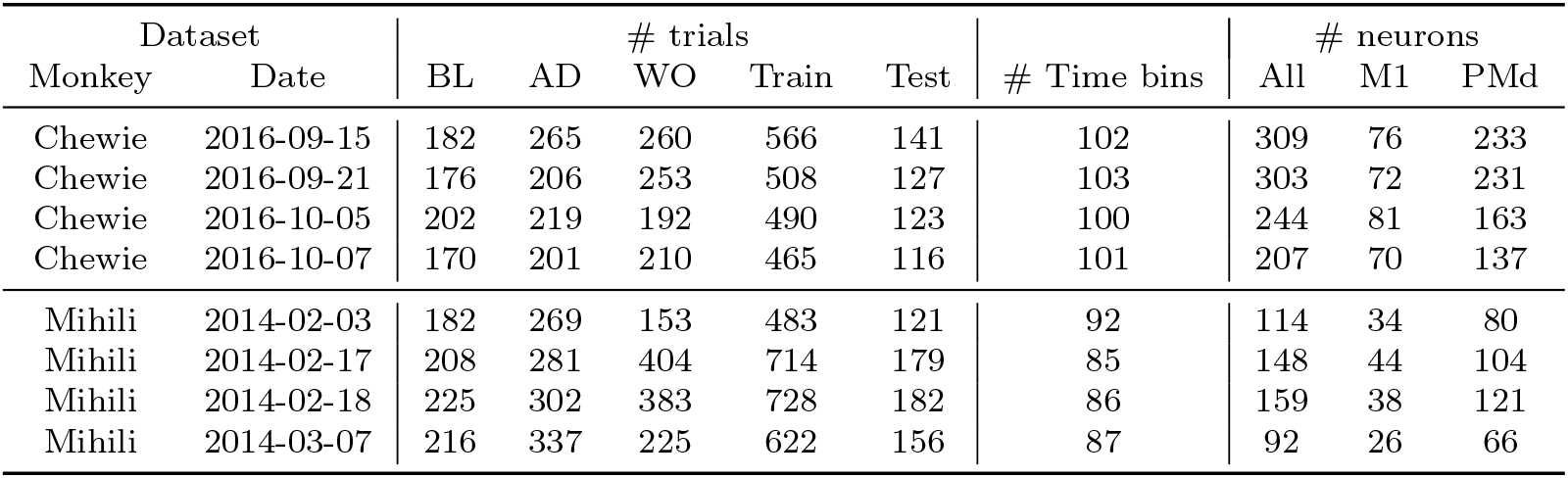
Datasets Information.

## Appendix C BAND captures additional behavioral variability encoded in small neural variability across multiple hand reaching tasks

We compared BAND to other generative models on three sets of Neural Latent Benchmark recordings from the motor cortex of monkeys performing different variants of hand reach tasks [36]. To compare the accuracy of the models in predicting behavior, we computed the coefficient of determination (R^2^) to measure the quality of the hand velocity reconstruction. To assess how well the models capture neural variability, we computed the Poisson likelihood and closely-related co-smoothing bits per second (co-bps [36]) as a measures of neural reconstruction quality.

We found that BAND outperformed all other non-ensemble models submitted to the benchmark in terms of the quality of hand velocity reconstruction (Fig. C3). This consistent high performance of BAND can not be trivially attributed to behavior supervision, since other models (CEBRA [39], MINT [38]) use behavior supervision too. The difference is, however, that BAND combines assumptions about neural dynamics on a lower-dimensional manifold (as in LFADS) with behavior supervision.

**Fig. C3.**
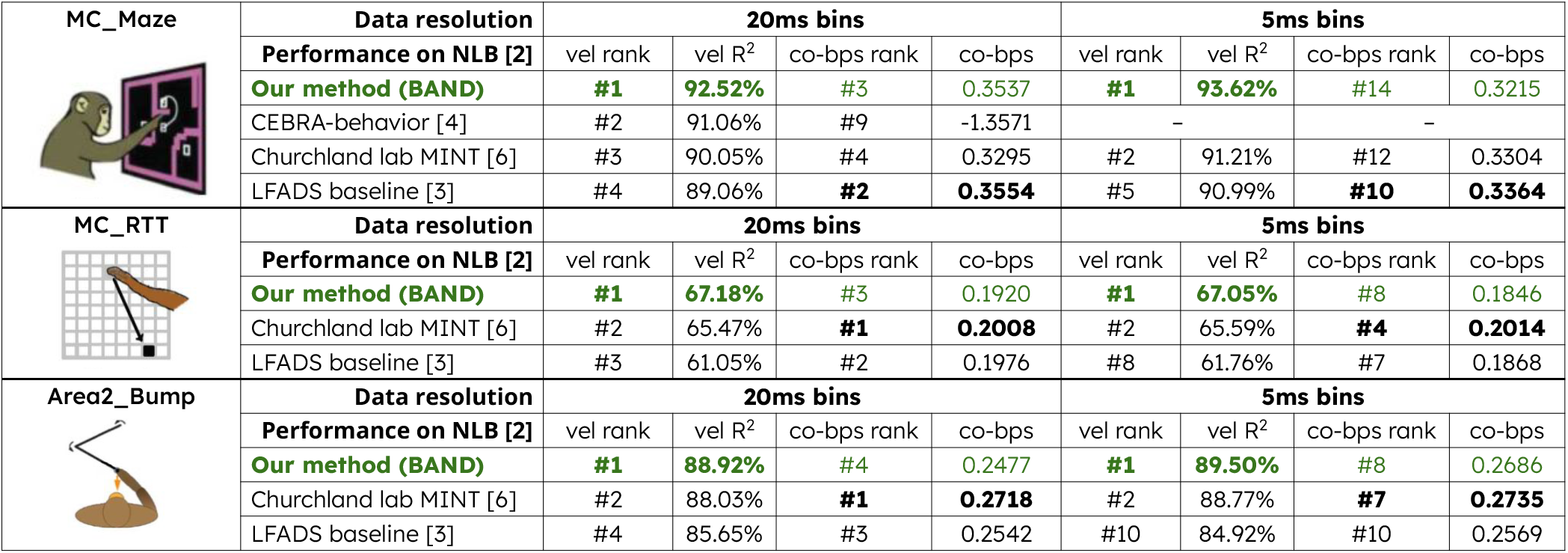
Validation on Neural Latents Benchmark [36]. The top 3 models with the highest behavior reconstruction quality are shown [5, 38]. Numbers in the table: hand velocity reconstruction (vel *R*^2^); neural reconstruction on held-out neurons (co-bps [36]); ranks show the position of the model on the [NLB leaderboard](https://neurallatents.github.io/) w.r.t. each metric (excluding ensemble models). BAND has the highest behavioral reconstruction among non-ensemble models but similar neural reconstruction to the LFADS baseline, across all NLB datasets.

Across all models in the benchmark, we found systematic differences in behavior prediction quality between the three hand-reaching tasks. For instance, in a fast-paced random reaching task (MC RTT, shown in Fig. C3), where one movement directly follows the other without any pre-movement delay periods, reconstruction quality is generally modest. In this case, the LFADS baseline reaches only *R*^2^ = 62%, while it achieves *R*^2^ = 89% in the center-out reach task (see MC Maze in Fig. C3), which has a clear trial structure. The MINT method from the Churchland lab reaches *R*^2^ = 65.5% due to its behavior-aware interpolation of latent states [38]. However, MINT still relies on the same AutoLFADS latents, which can limit performance due to the loss of behaviorally-relevant information in AutoLFADS latent space. At the same time, BAND had the largest relative improvement of behavior prediction on this random reach dataset compared to other two datasets (see Fig. C3), reaching *R*^2^ = 67%. This result suggests that our model is able to capture rapid, transient neural signals that are relevant to behavior, which can be missed by unsupervised latent state inference approaches like LFADS.

Notably, the improved behavior prediction in BAND is **not** related to improved neural activity reconstruction, compared to LFADS baseline. This was also observed in our previous semi-supervised model [12], and can be attributed to a slight reduction in the model’s ability to capture neural variability not related to behavior. This result suggests that the additional behavioral variability captured by BAND is encoded by small neural variability which has no significant impact on the reconstruction of neural firing.

Across all datasets, our method captured significant additional behavioral variability that corresponds to small neuronal variability.

### C.1 Adapting BAND for NLB challenge

Here we utilized the exact same version of LFADS that demonstrated strong performance on the leaderboard. We used a simple linear behavior decoder, architecturally identical to the Ridge regression model used by NLB for evaluating behavioral *R*^2^ (Fig. C4). While this version of the model was fine-tuned for the NLB challenge, its interpretability is limited, as well as its ability to capture behavioral feedback in neural activity due to the simplicity of the decoder.

**Fig. C4.**
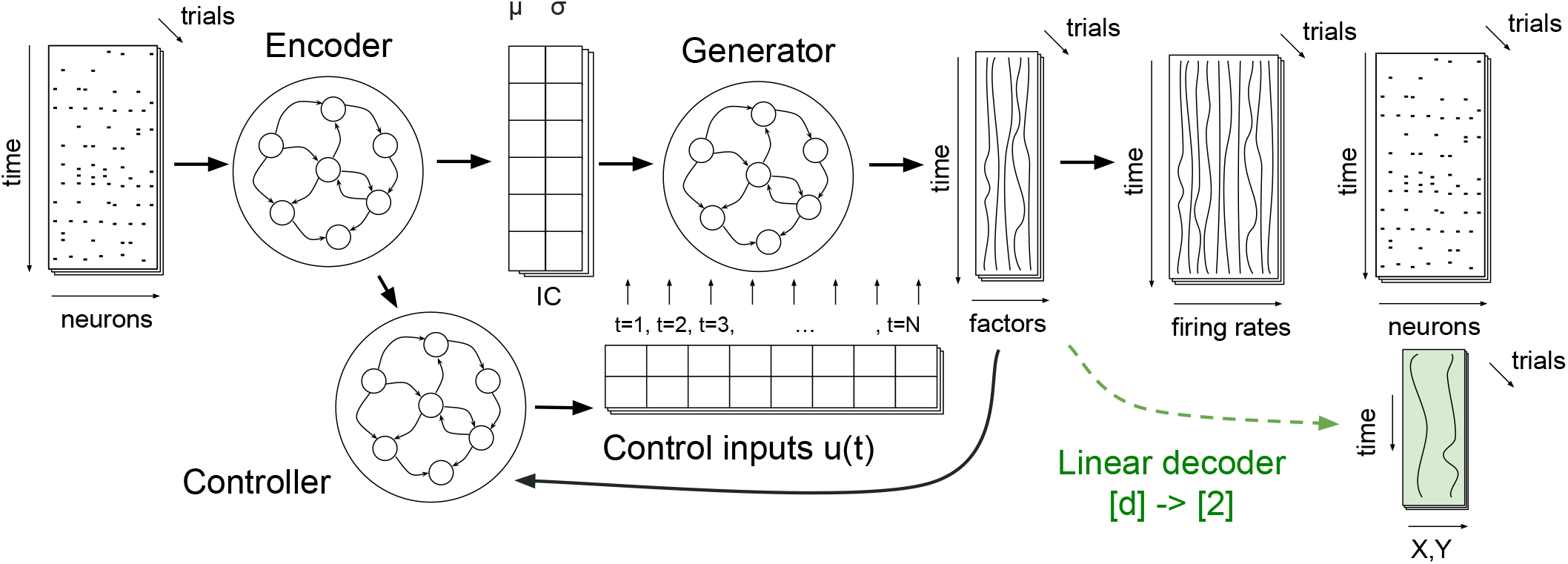
Acausal BAND archetecture with a fixed-lag behavior decoder for [Neural Latents Benchmark](https://neurallatents.github.io/) challenge.

We modified BAND decoder and objective in order to match the target metrics in NLB challenge. We used a standard linear decoder, instead of a sequence-to-sequence behavior decoder, to reproduce NLB decoding procedure. We also used the first two terms in BAND loss in (5) as a PBT objective for NLB challenge, with *θ* fixed to 10^−4^:

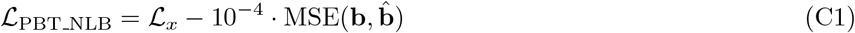

Both PBT losses in (7) and (C1) are equivalent up to constant coefficients before neural and behavioral terms. Note that there are multiple implementations of LFADS models in different frameworks: original in [Tensorflow 1](https://github.com/snel-repo/lfads_tf1), [Tensorflow 2](https://github.com/snel-repo/autolfads-tf2), and [PyTorch](https://github.com/arsedler9/lfads-torch). We used Tensorflow 2 version for the official NLB challenge, while the rest of the paper uses PyTorch version.

### Behavior supervision is essential for capturing the phase of movement correction in latent dynamics

We tested whether the information about hand velocity oscillations is preserved in latent dynamics models. These models summarize the activity of the neuronal population in a lower-dimensional set of latent factors. Different models follow different principles and objectives to identify latent variables [40]. Since we hypothesize that feedback-driven correction is encoded in a relatively small neural variability (Fig. 1), we expected that the methods that only rely on neural activity to identify latent variables, without any behavioral signal for supervision, will tend to discard movement corrections as noise. However, since movement corrections cause a substantial change in behavioral output, we expected that adding behavior supervision (Fig. 4) would ensure that the neural code for movement correction is included in the latent variables.

In this paper 5, we compared behavior reconstruction from the latent factors of two models: an unsupervised causal sequential autoencoder (CSAE) and our semi-supervised model (BAND). Although both models share an identical core architecture, they differ in their training methodology. BAND is trained end-to-end with a behavior decoder, whereas CSAE is trained without any behavioral supervision. To facilitate comparison, a linear sequence-to-sequence decoder was subsequently trained on the latent factors from the unsupervised CSAE model.

Latent factors in CSAE were able to capture the overall direction of movement, achieving *R*^2^ = 83% (Fig. C5c). This result is similar to a baseline model that predicts average hand trajectory knowing the true target direction (Fig. C5a, top). Decoded hand velocity from CSAE latent factors could also predict the 5 Hz oscillatory mode (Fig. C5c, middle), but not the phase of these oscillations (Fig. C5c, bottom; cosine similarity 0.27). This suggests that unsupervised nonlinear dynamics in CSAE could capture instructed reach direction, but could not explain and correctly time feedback-driven corrections.

**Fig. C5.**
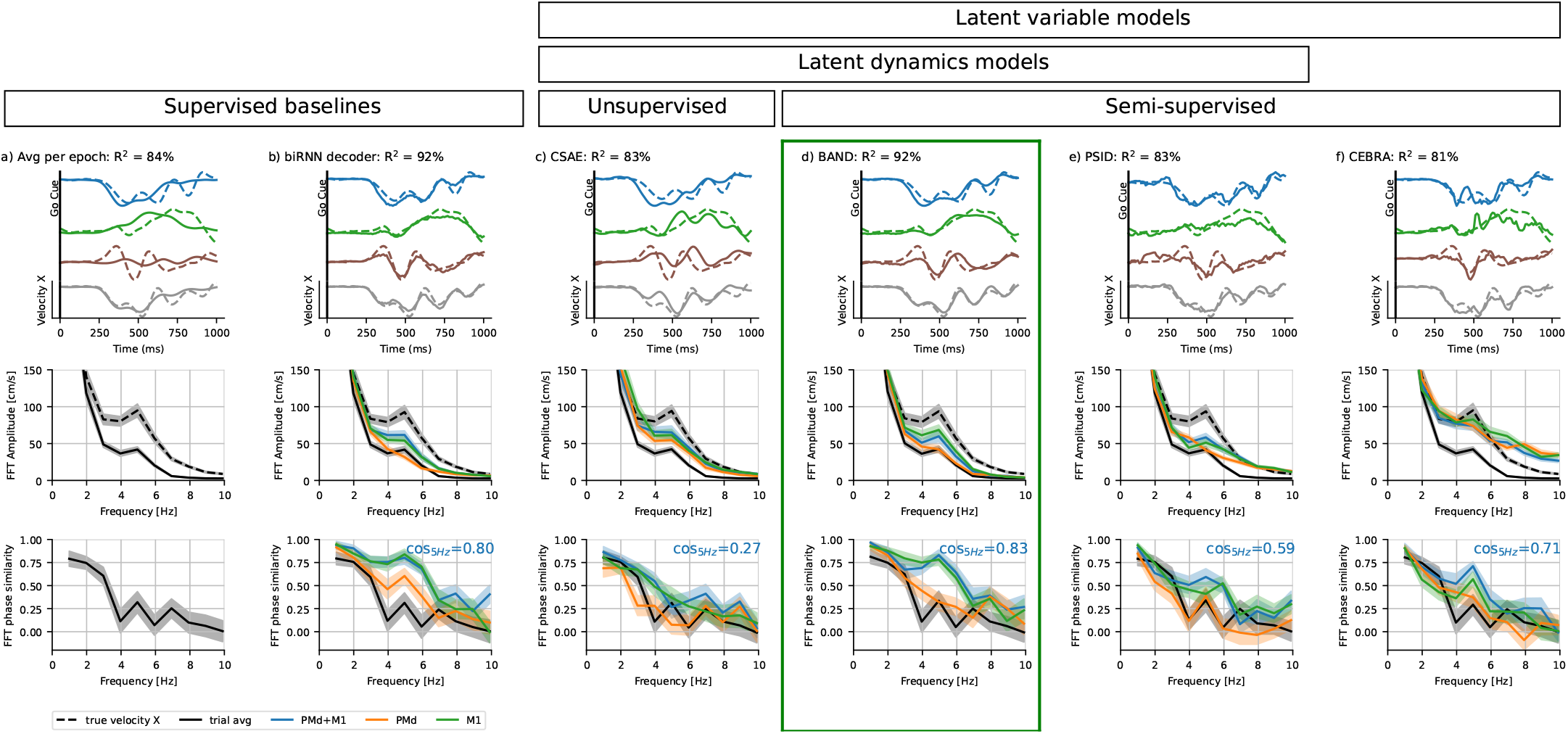
Model comparison shows that both nonlinear latent dynamics and behavior supervision are required for capturing hand velocity. Top row: hand velocity in example AD trials (solid line – model prediction from PMd+M1, dashed – ground truth); middle row: a Fourier spectrum of hand velocity predictions (from both or either brain areas), indicating whether higher amplitude of 5 Hz oscillations is correctly captured; bottom row: cosine similarity between Fourier modes of true velocity vs predicted velocity; cosine similarity at 5 Hz indicates whether the phase of oscillations is captured; **a)** An average velocity towards the reach target (average across all trials with the same reach target within AD epoch); **b)** Velocity predicted by a supervised bi-directional RNN decoder, demonstrating that hand velocity oscillations are decodable from neural activity; **c)** Velocity predicted using ridge regression from CSAE model with a 4-dimensional controller and 100 latent factors; **d)** Velocity predicted by BAND model with 100 factors and all hyperparameters matched to a CSAE model; **e)** Velocity predicted using a kNN decoder from CEBRA embedding; **f)** Velocity predicted by PSID model (a linear Kalman filter with neural and behavioral observations). Decoding here was performed on both PMd and M1 using all epochs, and results evaluated on adaptation epoch; predictions for all reach directions and individual brain areas can be found in Fig. H13)

Unlike CSAE, BAND achieved high accuracy in behavior reconstruction, *R*^2^ = 92%, matching the biRNN decoder performance in overall hand trajectory reconstruction (Fig. C5d, top) and capturing the phase of the oscillations (Fig. C5d, bottom). BAND can summarize both movement plan and correction in the latent dynamics without a significant loss of neural reconstruction performance compared to CSAE (0.264 vs. 0.259 bits / second for the models shown here; standard deviation for cross-validated neural reconstruction is 0.004 bits / second). The fact that a CSAE model optimized for neural reconstruction quality discards these feedback-driven motor corrections suggests that they are encoded in low-variance patterns of neural activity.

#### Alternative semi-supervised models with fixed-lag decoders fail to capture oscillations

We tested whether non-linear dynamics was essential for achieving a high behavior reconstruction for center-out reaching. In theory, the overall hand reaching plan can be represented as a rotation in a latent space [41], which is a linear dynamics mode. The movement corrections to a force field perturbation, as illustrated in Fig. 2, mostly correspond to a 5 Hz oscillation, which is also a linear mode once it is switched on in perturbed trials. Therefore, both important dynamical modes can be represented with linear dynamics, as long as these modes can be switched on and off.

We therefore compared our results with another semi-supervised, yet linear, dynamical model, Preferential sub-space identification (PSID [20]). PSID is based on a linear Kalman smoother, which summarizes neural activity into linear combinations of latent factors with linear dynamics. As a Kalman smoother, PSID combines linear dynamic predictions with incoming evidence from the data, which can, in principle, account for nonlinear switches between dynamic modes (e.g. preparation to movement). An additional feature of PSID is that the latent space is split into two: a behaviorally relevant subspace that dissociates and prioritizes behaviorally relevant dynamics, and a behaviorally irrelevant subspace. The behaviorally relevant subspace is assumed to be temporally aligned with the neural activity (possibly, with a fixed lag), which is more restrictive than the linear sequence-to-sequence decoding in BAND.

We first assessed the eigenvalues of the learned dynamics in PSID and found the minimal number of factors that gives rise to latent neural oscillations with a period of 200 *±* 100 ms (5 Hz). We found that PSID needs at least 6 behaviorally relevant factors. PSID with only behaviorally irrelevant factors (which is, essentially, an unsupervised Kalman smoother of neural activity) needs *>*17 factors. If unsupervised PSID is trained on perturbed trials only, then it needs at least 9 factors to predict oscillations. This result suggests that oscillations are encoded in high-order components of a linear state-space model.

However, these 5 Hz oscillations in PSID hand velocity predictions appeared to be out of phase. We tested the behavior reconstruction performance of the PSID model with 100 factors (same as LFADS/BAND). We optimized the number of behaviorally relevant factors to ensure the most accurate decoding of movement in this dataset, resulting in 40 relevant and 60 irrelevant factors. Despite capturing some neural oscillations with the 5 Hz frequency, the overall behavior reconstruction quality was relatively low (*R*^2^ = 83%, Fig. C5e, top), comparable to condition-averaged model (*R*^2^ = 84%, Fig. C5a, top). In general across the adaptation trials, the oscillations were not well-captured, with both the amplitude and the phase of the 5 Hz component poorly predicted (Fig. C5e, middle-bottom rows).

Finally, we tested whether behavior supervision alone, without latent dynamics, is sufficient to obtain a latent representation of movement plan and correction. We tested this hypothesis using a recently proposed supervised embedding method CEBRA [39]. CEBRA accounted for some motor corrections, but also added high-frequency features to velocity predictions that are absent from the original signal (Fig. C5f, middle row). CEBRA failed to reproduce a distinct peak in amplitude of 5 Hz oscillation frequency, yet captured the phase of oscillations more accurately than phases in other frequency bands. However, the accuracy of reproducing the phase was still low: the cosine similarity between the true phase and the predicted oscillation phase was 0.66: comparable to PSID (0.61), but substantially lower than that of the biRNN decoder (0.80) or BAND (0.83). This suggests that behavior supervision alone, without explicitly modeling neural dynamics, is insufficient for capturing motor corrections.

Throughout the paper, we evaluate model performance in behavior reconstruction with the *R*^2^ computed on the 2D velocity. However, visualizations and FFT analysis often display only one of the velocity components, for the sake of simplicity. For instance, Figure C5 shows the prediction for velocity component X. Here we show that the benchmarking results for different types of models still holds if evaluated on another component (Fig. C6) or the norm of the velocity (Fig. C7).

**Fig. C6.**
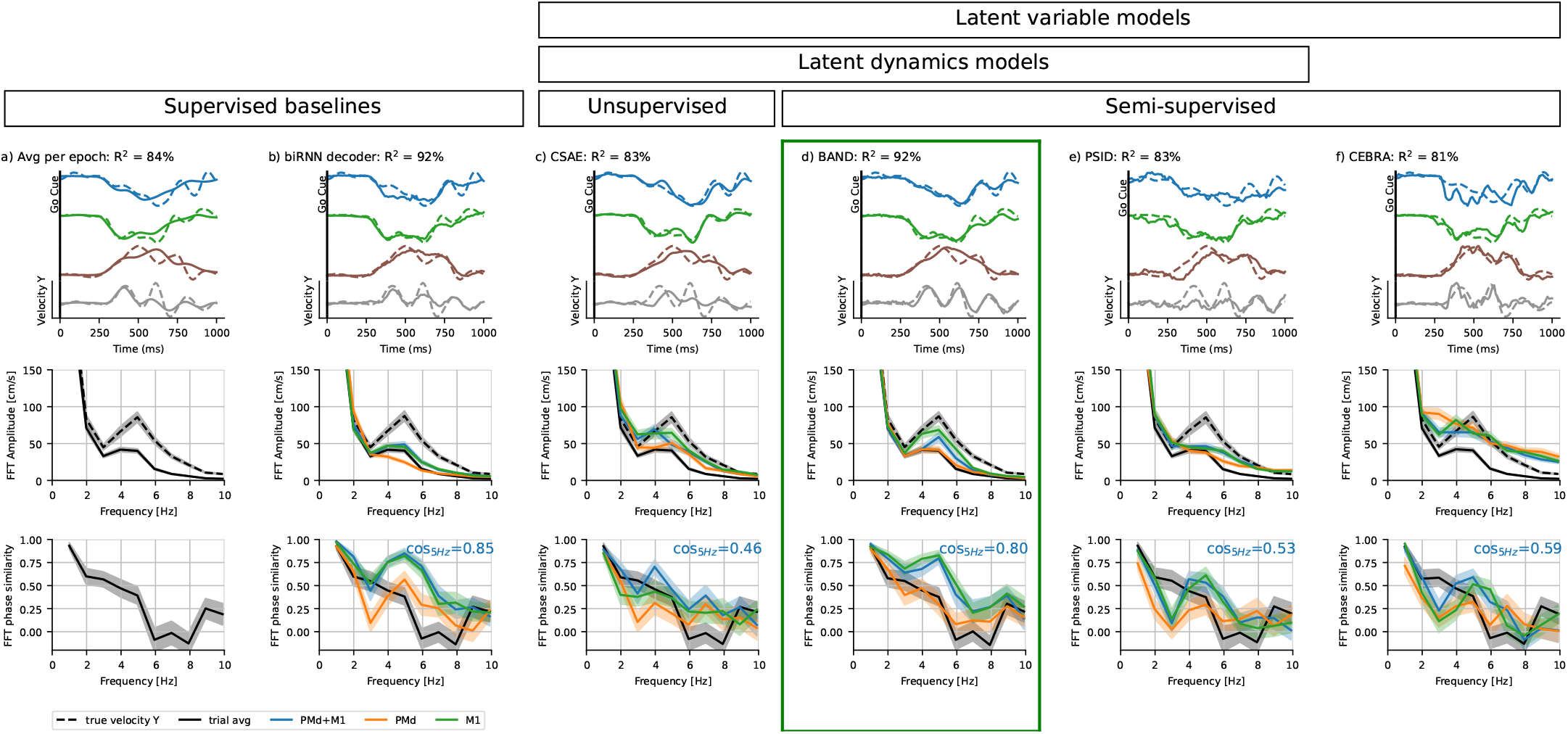
Same as Figure C5, but with velocity component Y.

**Fig. C7.**
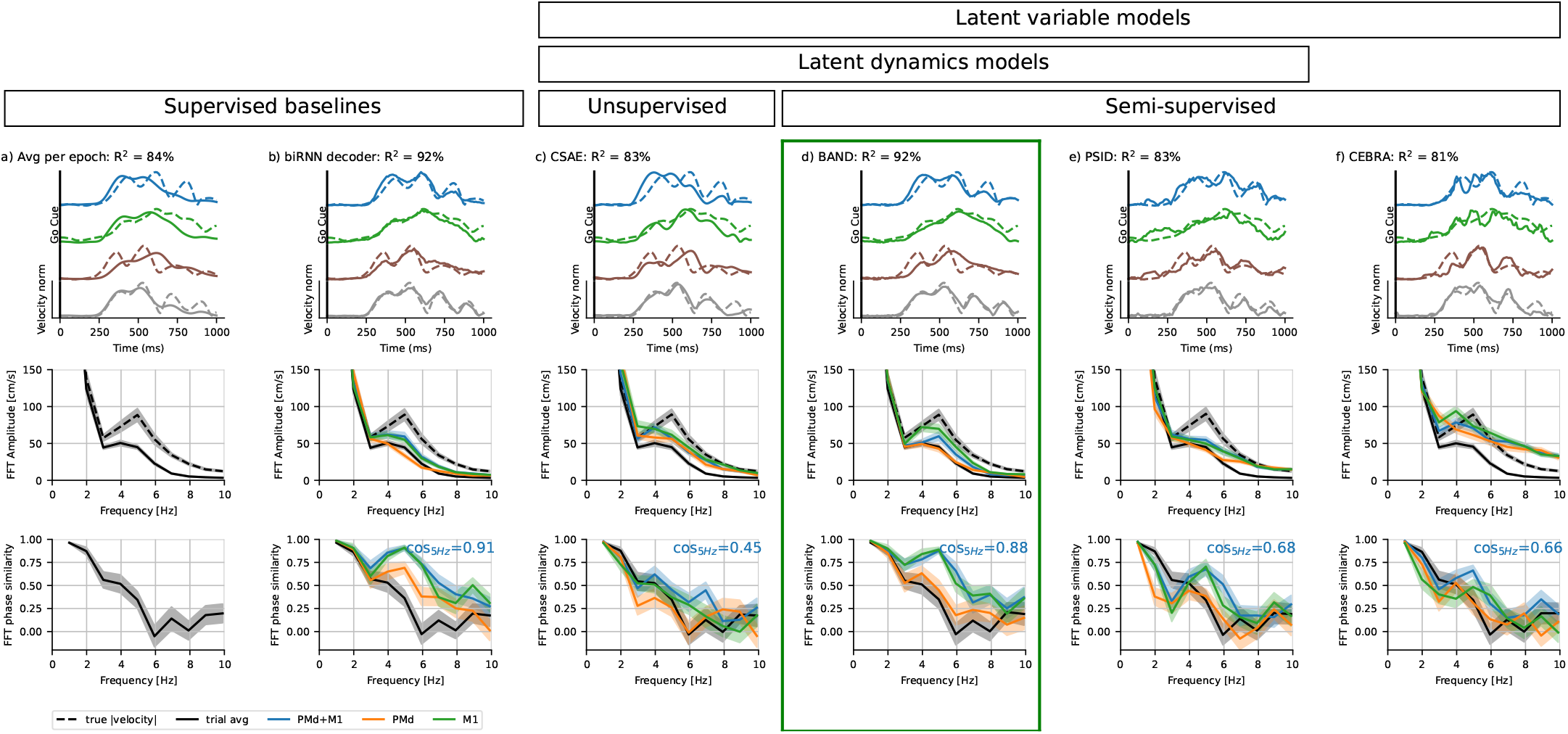
Same as Figure C5, but with the norm of hand velocity.

### C.2 Model parameters for this benchmark

#### CEBRA and a kNN decoder

We used a CEBRA [39] offset10-model to obtain the 100-dimensional embeddings of neural activity. We used batch size of 512, learning rate 10^−4^ and temperature 1, and trained the model for 10000 iterations with the cosine similarity as the distance metric. We used velocity as a continuous label for CEBRA embeddings.

We then used kNN regression to predict hand velocity, and the number of neighbors k was searched over the range [1, 2500] (same as in the original CEBRA[39]). We fine-tuned the number of neighbors for each dataset based on cosine similarity.

#### PSID decoder

We used PSID model [20] with 100 factors. We optimized the dimensionality of behaviorally-relevant subspace to achieve the highest behavior decoding performance (*R*^2^), which resulted in 40 factors, leaving 60 behaviorally-irrelevant factors.

To make behavior predictions, we used the fixed-time linear behavior decoder from the PSID model. This decoder takes the latent state at a given moment in time and predicts behavioral state at the same time.

## APPENDIX D Testing autonomous models (no controller trained)

Here we test whether autonomous models, i.e. without a controller, can learn latent factors predictive of hand velocity oscillations. This test demonstrates that the **causal** restrictions applied on our model prevent the model from compressing all the information about the trial into initial condition – a possible solution in original acausal LFADS, which lacks interpretability.

None of the models could predict the phase of oscillations correctly, even though acausal models had access to the future spiking data (bottom, Fig. D8). However, tuning all hyperparameters in the acausal models, as it is done in AutoLFADS [37], results in poorly regularized models with still create 5 Hz oscillations (middle row, Fig. D8), but fail to synchronize the phase. Adding causality prevents autonomous models from creating spurious oscillations.

**Fig. D8.**
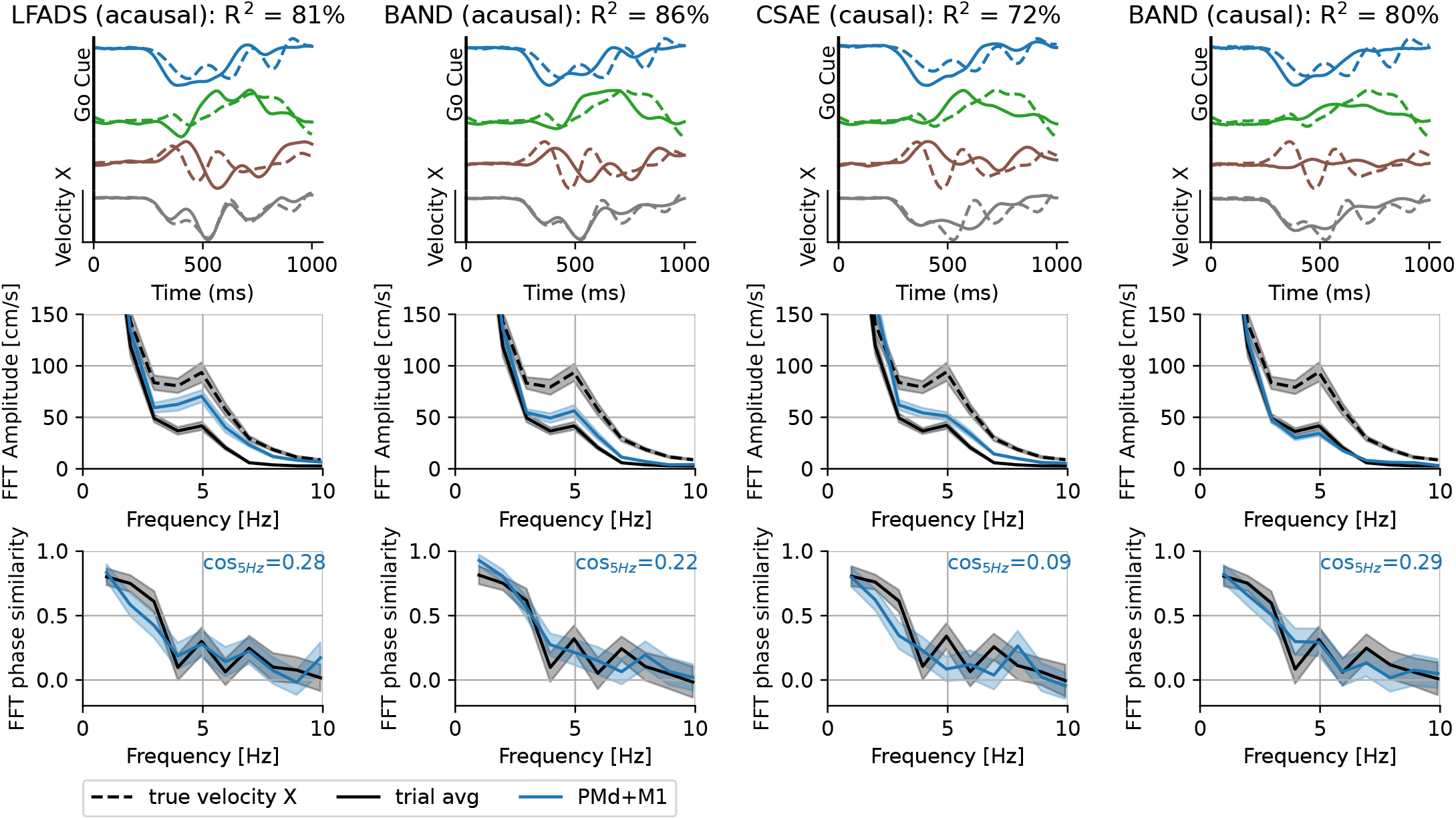
Model comparison for **autonomous** LFADS/BAND models, i.e. with controller removed from the model architecture.

## APPENDIX E Control inputs in BAND cause only a transient deviation from an autonomous trajectory

Inputs inferred via BAND, despite their transient nature, are not sparse in time; rather, they continually modulate the dynamics. By isolating individual inputs, we show that each input induces transient deviations from an autonomous trajectory without diverging from the long-term trajectory.

By masking inputs in all but one time bin, we assessed how single-point perturbations affect latent and behavioral trajectories (Fig. E9). The resulting deviations (measured as a square distance) were transient, lasting less than 200 ms (see autocorrelogram half-width).

**Fig. E9.**
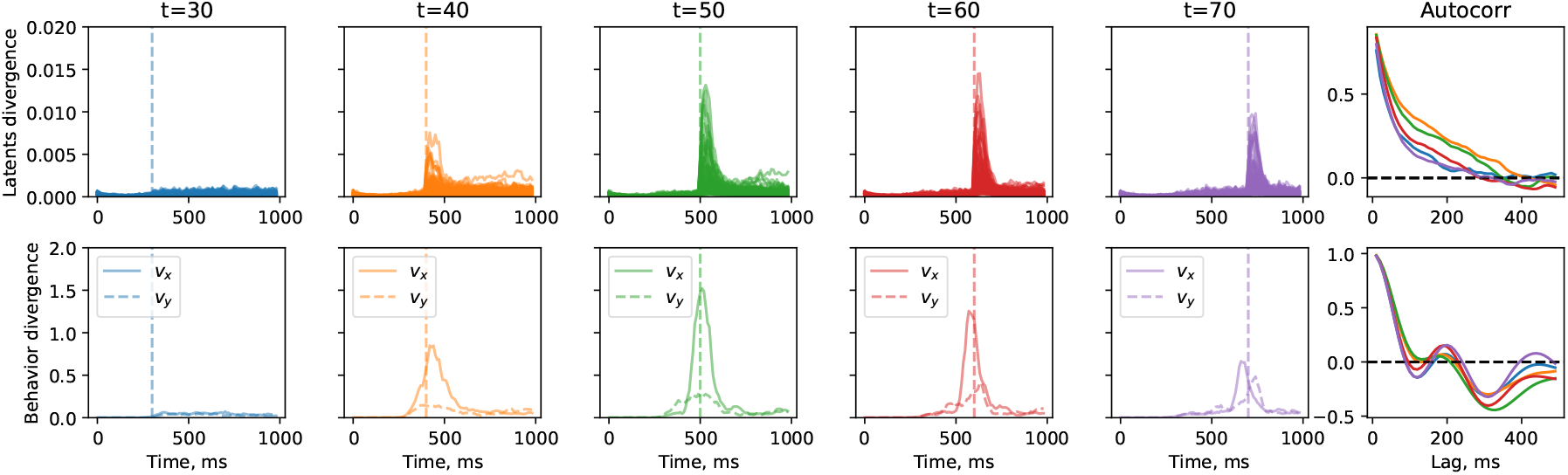
Deviation from the autonomous trajectory in the latent space and behavioral output space. Each line in the first five columns shows the trial-average residual along one of the dimensions. The last column shows autocorrelograms for the residuals evoked by perturbations.

## APPENDIX F Control inputs in BAND correlate with movement speed

**Fig. F10.**
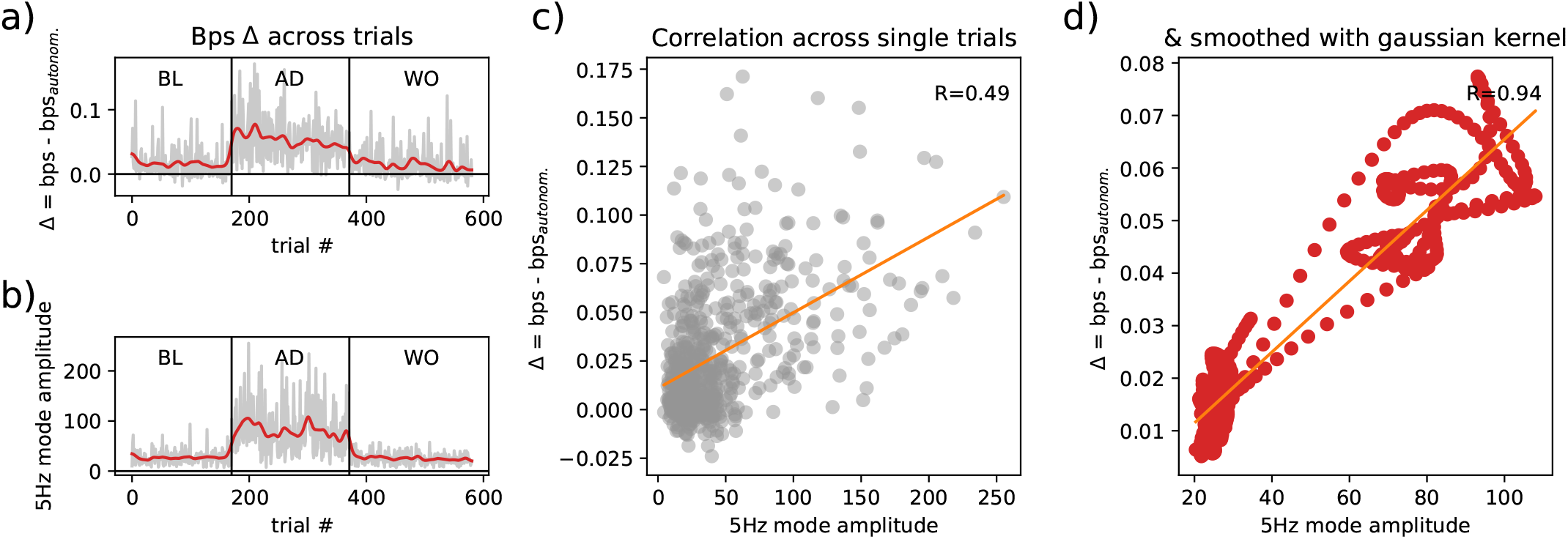
The controller adds oscillations to neural factors. a) BAND controller explains more neural variability (bits / second) in adaptation trials than in other epochs without perturbation; b) amplitude of the 5 Hz hand velocity oscillations (reproduces Fig. 2e, left); c) Correlation between single trial neural variability explained by controller (bps) and amplitude of the 5 Hz hand velocity oscillation mode (R=0.49); d) Correlation between across-trials smoothed trends (gaussian kernel s.d.=5 trials) of neural variability explained by controller (bps) and amplitude of the 5 Hz hand velocity oscillation mode (R=0.94);

**Fig. F11.**
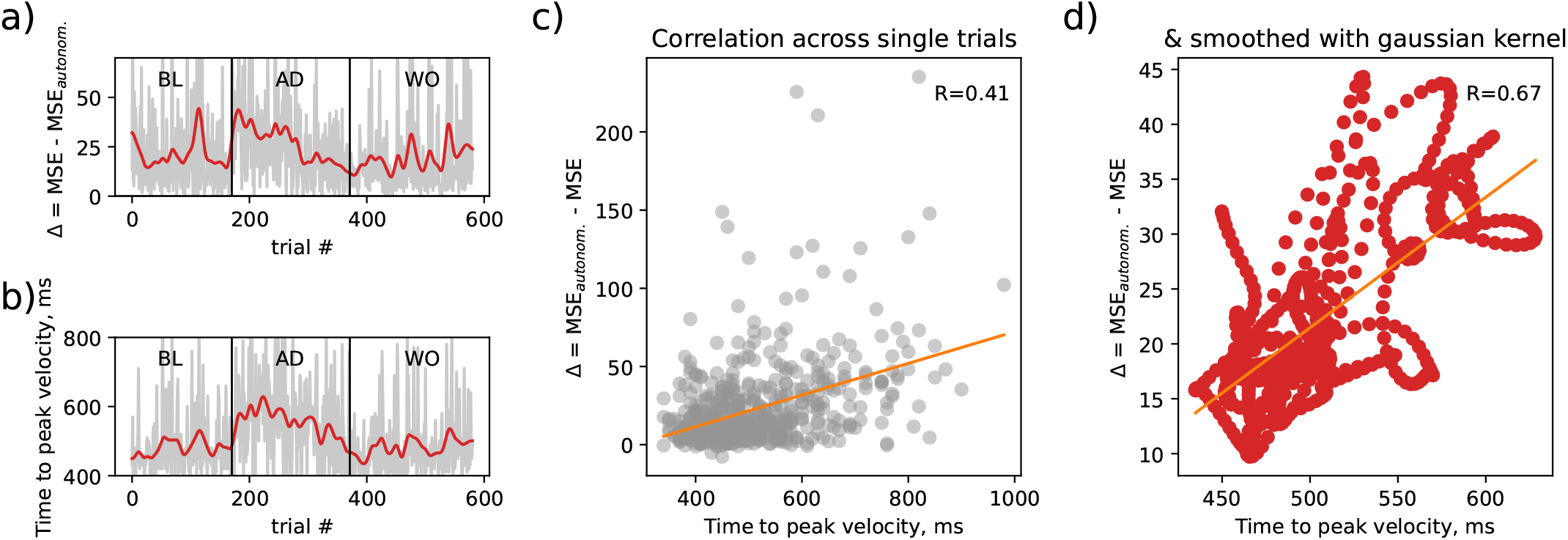
The controller contributes more to behavior prediction in the trials when reaching max velocity was delayed. a) Drop in mean-squared error due to controller ablation in BAND; b) Time from the start of the trial (movement onset - 250 ms) to reaching peak velocity; c) Correlation between single trial ΔMSE and time to reach peak velocity (R=0.41); d) Correlation between across-trials smoothed trends (gaussian kernel s.d.=5 trials) of ΔMSE and time to reach peak velocity (R=0.67);

## APPENDIX G Visualization of BAND latents

**Fig. G12.**
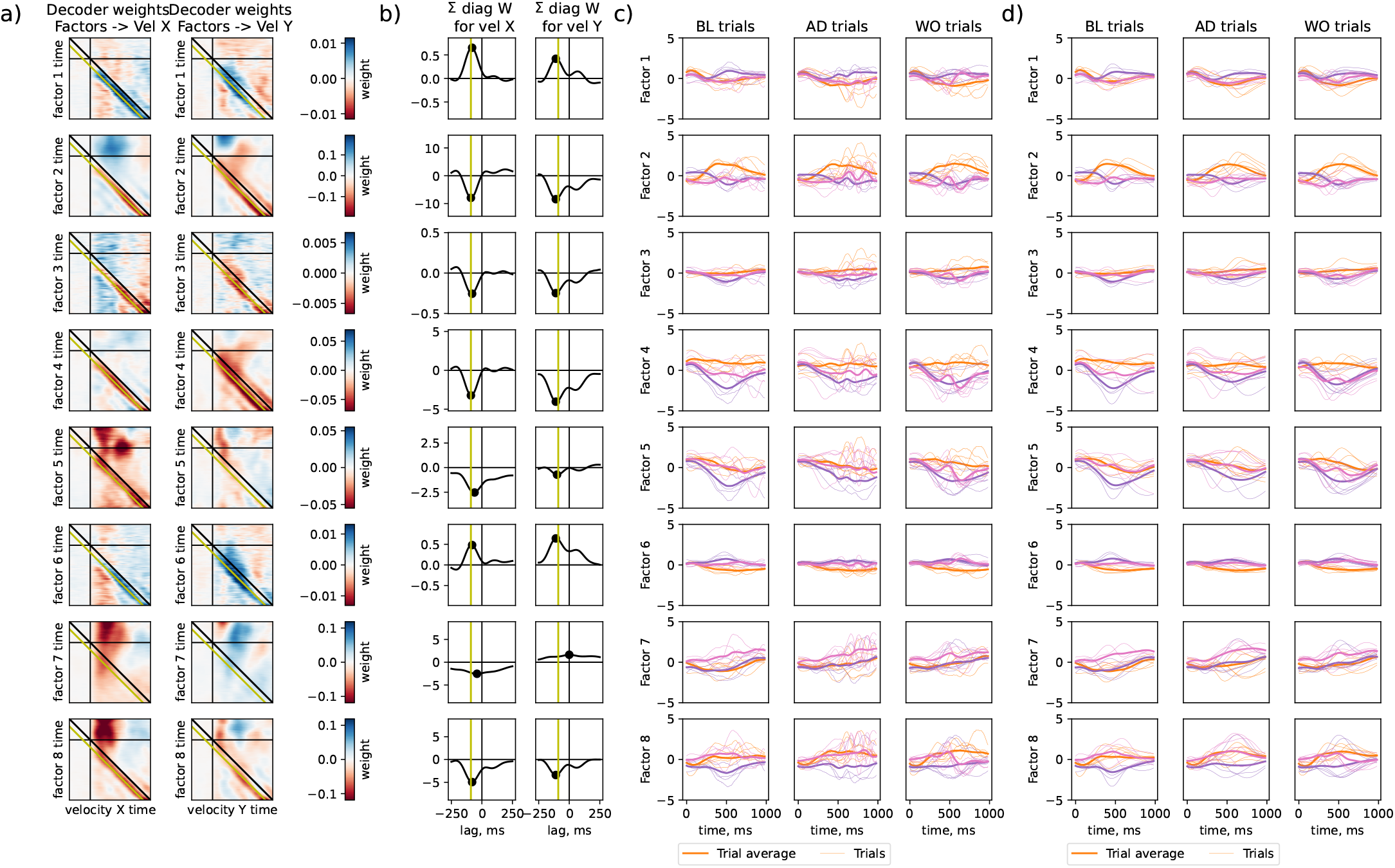
. Behavior decoder weights and factors for a 8-factor BAND model with a causal controller.

## APPENDIX H Decoding performances (R^2^ and cosine similarity) and trajectory divergence for different areas, methods, and sessions

We applied Fourier decomposition to hand velocity predictions from biRNN decoder (Fig. H13), LFADS (Fig. H14) and BAND (Fig. H16).

**Fig. H13.**
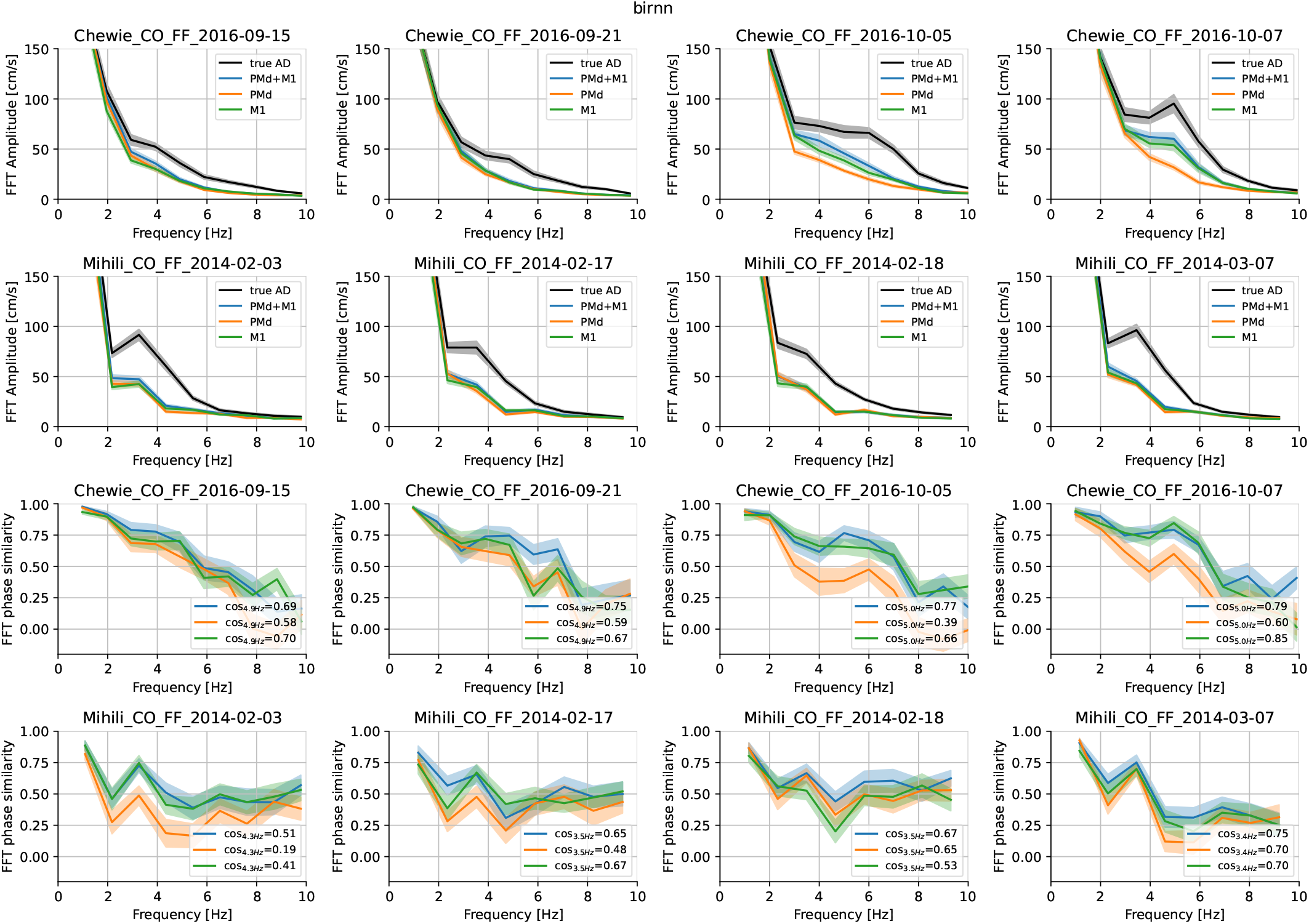
Fourier spectrum for velocity predictions based on biRNN decoder. In the sessions where hand velocity oscillations are decodable, they are decodable from M1 and not PMd. Note, that the number of M1 neurons recorded in Monkey M is considerably lower then in Monkey C. Brain areas used in the decoding are color-coded (blue: PMd+M1, orange: PMd, green: M1).

**Fig. H14.**
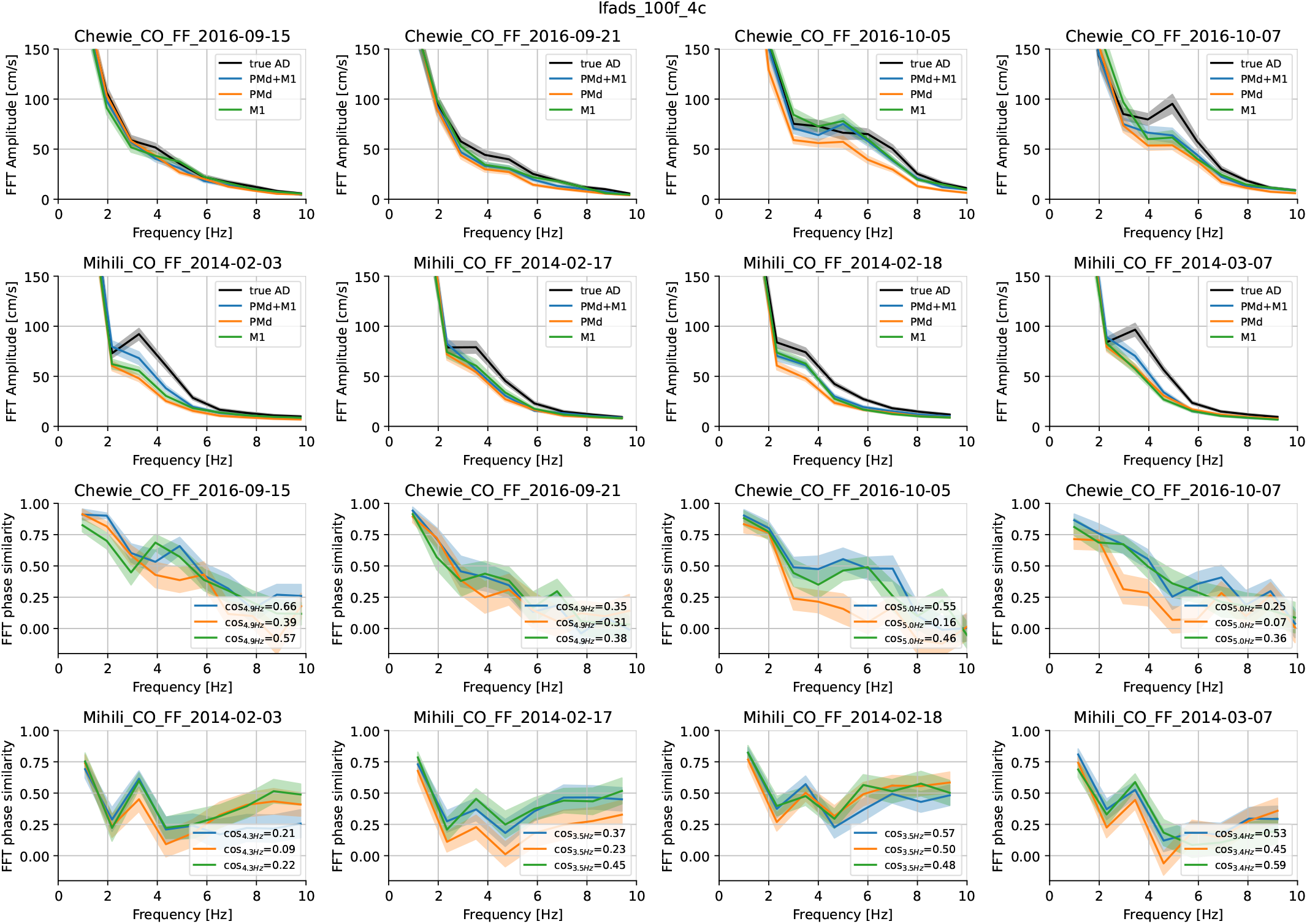
Fourier spectrum for velocity predictions based on CSAE models with 100 factors.

**Fig. H15.**
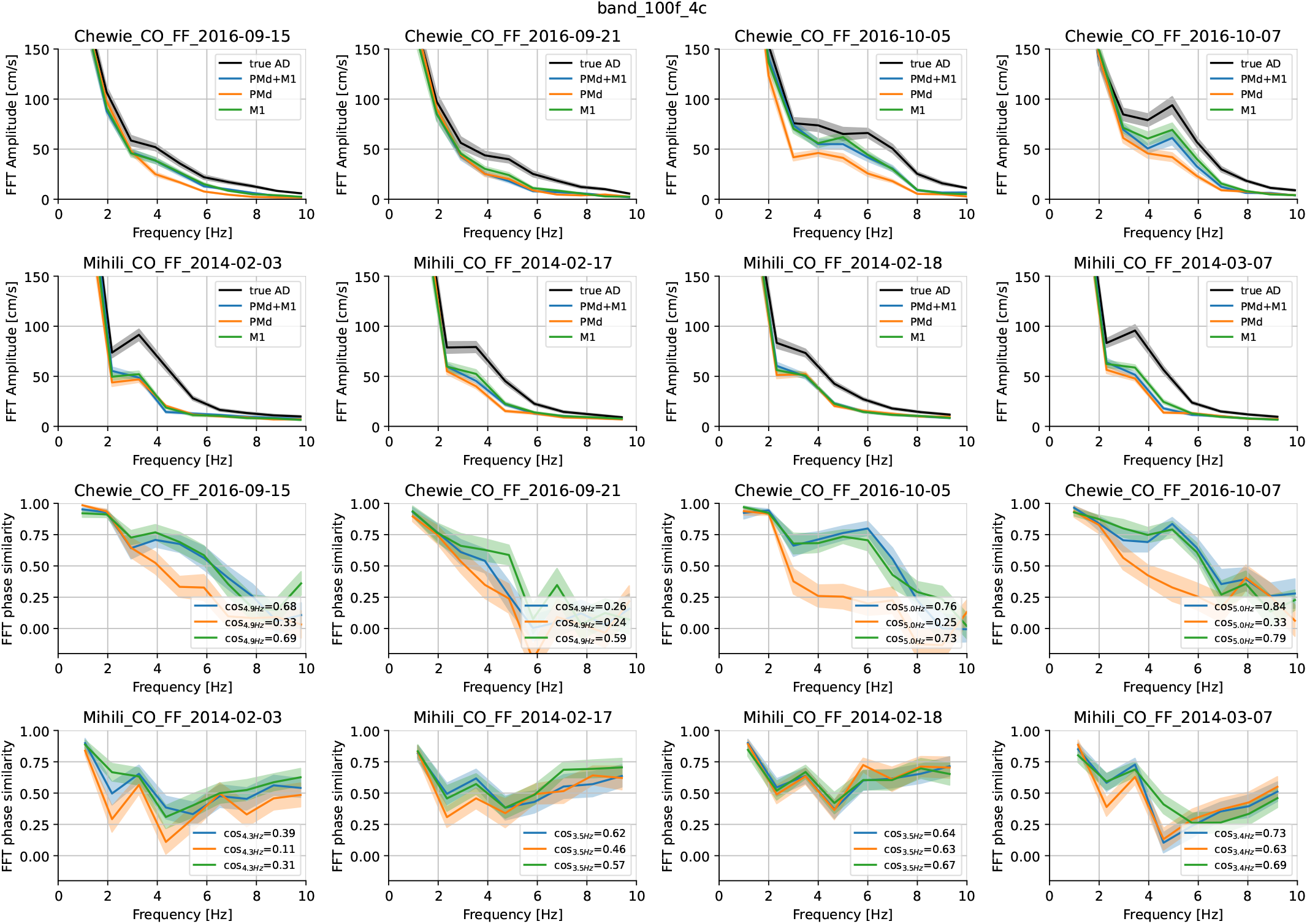
Fourier spectrum for velocity predictions based on BAND models with 100 factors.

**Fig. H16.**
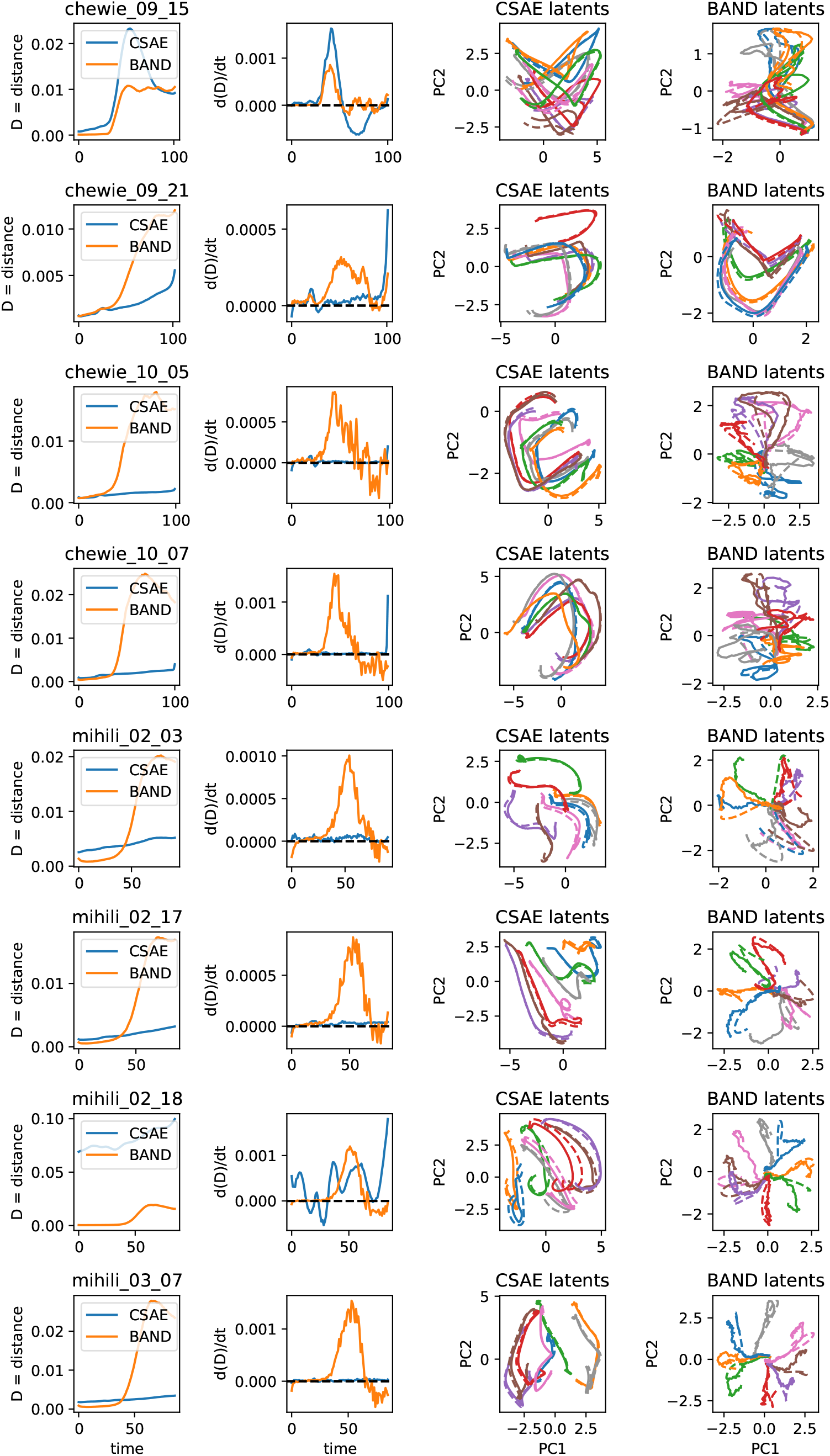
Latent trajectory divergence for controlled vs. autonomous latent trajectories in CSAE / BAND across animals and sessions. All neurons (M1 and PMd) were used in training these models, and all datasets were aligned to go cue. BAND shows consistent transient trajectory divergence during movement (orange traces), which then stabilizes (d(D))/dt=0) or even begins to converge (d(D)/dt*<*0). CSAE demonstrates a variety of behaviors (blue traces), with most models slightly diverging from the autonomous trajectories towards the end.

**Table H3.**
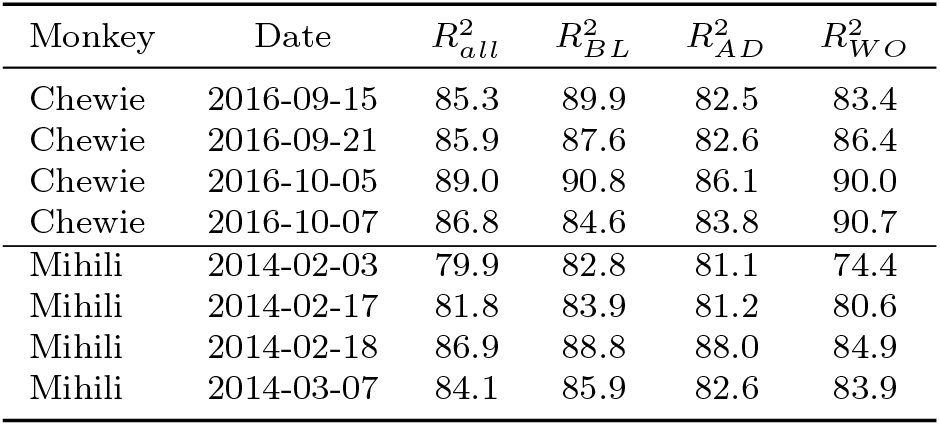
Variance explained by an average hand velocity towards a given target (*R*^2^, %)

**Table H4.**
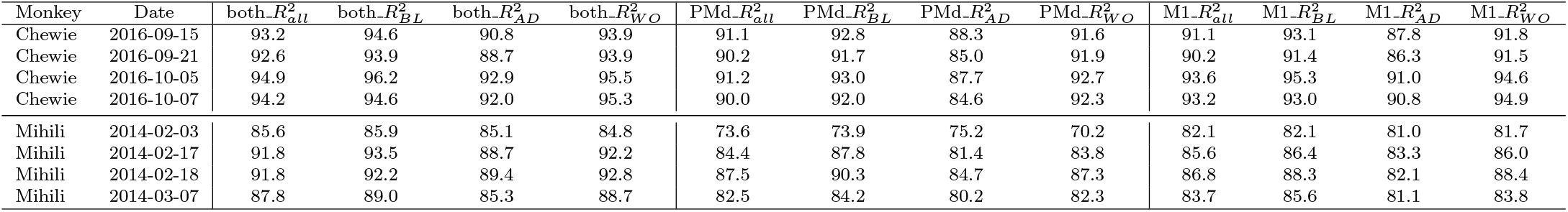
Variance explained by a biRNN decoder given neural recordings from M1, PMd, or both areas together (*R*^2^, %). Red indicates cases in which biRNN decoding captured less variance than a model that only predicts reach direction (i.e. *R*^2^ is below that in Table H3).

**Table H5.**
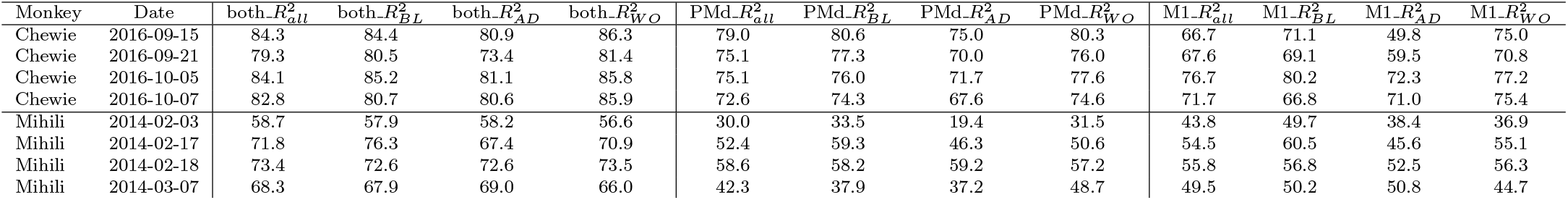
Variance explained by a CEBRA + kNN decoder given neural recordings from M1, PMd, or both areas together (*R*^2^, %). One green value indicates cases in which CEBRA explained more variance than the average hand trajectory (i.e. Table. H3).

**Table H6.**
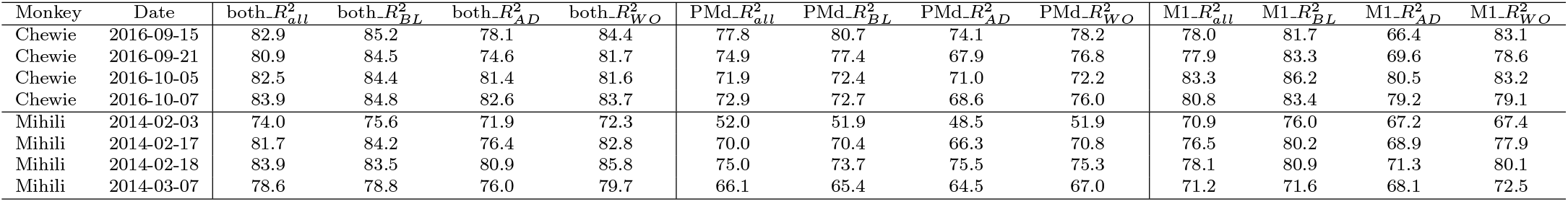
Variance explained by a PSID decoder given neural recordings from M1, PMd, or both areas together (*R*^2^, %). Green values indicate cases in which PSID explained more variance than the average hand trajectory (i.e. Table. H3). Note, that PSID never explained uninstructed variance in perturbed trials.

## References

[1] Churchland, M. M., Afshar, A. & Shenoy, K. V. A central source of movement variability. Neuron 52, 1085–1096 (2006).

[2] Vyas, S., Golub, M. D., Sussillo, D. & Shenoy, K. V. Computation through neural population dynamics. Annual review of neuroscience 43, 249–275 (2020).

[3] Kao, T.-C., Sadabadi, M. S. & Hennequin, G. Optimal anticipatory control as a theory of motor preparation: A thalamo-cortical circuit model. Neuron 109, 1567–1581 (2021).

[4] Schimel, M., Kao, T.-C., Jensen, K. T. & Hennequin, G. ilqr-vae: control-based learning of input-driven dynamics with applications to neural data. bioRxiv 2021–10 (2021).

[5] Pandarinath, C. et al. Inferring single-trial neural population dynamics using sequential auto-encoders. Nature methods 15, 805–815 (2018).

[6] Perich, M. G., Gallego, J. A. & Miller, L. E. A neural population mechanism for rapid learning. Neuron 100, 964–976 (2018).

[7] Gandolfo, F., Li, C.-S., Benda, B., Schioppa, C. P. & Bizzi, E. Cortical correlates of learning in monkeys adapting to a new dynamical environment. Proceedings of the National Academy of Sciences 97, 2259–2263 (2000).

[8] Li, C.-S. R., Padoa-Schioppa, C. & Bizzi, E. Neuronal correlates of motor performance and motor learning in the primary motor cortex of monkeys adapting to an external force field. Neuron 30, 593–607 (2001).

[9] Arce, F., Novick, I., Mandelblat-Cerf, Y. & Vaadia, E. Neuronal correlates of memory formation in motor cortex after adaptation to force field. Journal of Neuroscience 30, 9189–9198 (2010).

[10] Sun, X. et al. Cortical preparatory activity indexes learned motor memories. Nature 602, 274–279 (2022).

[11] Dooley, J. C. & Blumberg, M. S. Developmental’awakening’of primary motor cortex to the sensory consequences of movement. elife 7, e41841 (2018).

[12] Hurwitz, C. et al. Targeted neural dynamical modeling. NeurIPS 34, 29379–29392 (2021).

[13] Georgopoulos, A., Kalaska, J., Caminiti, R. & Massey, J. Interruption of motor cortical discharge subserving aimed arm movements. Experimental Brain Research 49, 327–340 (1983).

[14] Pruszynski, J. A. et al. Primary motor cortex underlies multi-joint integration for fast feedback control. Nature 478, 387–390 (2011).

[15] Scott, S. H. Optimal feedback control and the neural basis of volitional motor control. Nature Reviews Neuroscience 5, 532–545 (2004).

[16] Versteeg, C. & Miller, L. Dynamical feedback control: motor cortex as an optimal feedback controller based on neural dynamics (2022).

[17] Gurnani, H., Liu, W. & Brunton, B. W. Feedback control of recurrent dynamics constrains learning timescales during motor adaptation. bioRxiv 2024–05 (2024).

[18] Michaels, J. A. et al. Sensory expectations shape neural population dynamics in motor circuits. Nature 1–10 (2025).

[19] Perich, M. G., Narain, D. & Gallego, J. A. A neural manifold view of the brain. Nature Neuroscience 1–16 (2025).

[20] Sani, O. G., Abbaspourazad, H., Wong, Y. T., Pesaran, B. & Shanechi, M. M. Modeling behaviorally relevant neural dynamics enabled by preferential subspace identification. Nature Neuroscience 24, 140–149 (2021).

[21] Sani, O. G., Pesaran, B. & Shanechi, M. M. Dissociative and prioritized modeling of behaviorally relevant neural dynamics using recurrent neural networks. Nature neuroscience 27, 2033–2045 (2024).

[22] Feulner, B., Perich, M. G., Miller, L. E., Clopath, C. & Gallego, J. A. Feedback-based motor control can guide plasticity and drive rapid learning. bioRxiv (2022).

[23] Feulner, B., Perich, M. G., Miller, L. E., Clopath, C. & Gallego, J. A. A neural implementation model of feedback-based motor learning. Nature Communications 16, 1805 (2025).

[24] Perich, M. G. et al. Motor cortical dynamics are shaped by multiple distinct subspaces during naturalistic behavior. BioRxiv 2020–07 (2020).

[25] Cisek, P. & Green, A. M. Toward a neuroscience of natural behavior. Current opinion in Neurobiology 86, 102859 (2024).

[26] Ye, J. & Pandarinath, C. Representation learning for neural population activity with neural data transformers. arXiv preprint arXiv:2108.01210 (2021).

[27] Le, T. & Shlizerman, E. Stndt: Modeling neural population activity with spatiotemporal transformers. Advances in Neural Information Processing Systems 35, 17926–17939 (2022).

[28] Azabou, M. et al. A unified, scalable framework for neural population decoding. Advances in Neural Information Processing Systems 36, 44937–44956 (2023).

[29] Dowling, M., Zhao, Y. & Park, M. exponential family dynamical systems (xfads): Large-scale nonlinear gaussian state-space modeling. Advances in Neural Information Processing Systems 37, 13458–13488 (2024).

[30] Ali, Y. H. et al. Brand: a platform for closed-loop experiments with deep network models. Journal of Neural Engineering 21, 026046 (2024).

[31] Safaie, M. et al. Preserved neural dynamics across animals performing similar behaviour. Nature 623, 765–771 (2023).

[32] Ye, J., Collinger, J., Wehbe, L. & Gaunt, R. Neural data transformer 2: multi-context pretraining for neural spiking activity. bioRxiv 2023–09 (2023).

[33] Zhang, Y. et al. Towards a” universal translator” for neural dynamics at single-cell, single-spike resolution. Advances in Neural Information Processing Systems 37, 80495–80521 (2025).

[34] Zhang, Y. et al. Exploiting correlations across trials and behavioral sessions to improve neural decoding. bioRxiv (2024).

[35] Maeda, R. S., Kersten, R. & Pruszynski, J. A. Shared internal models for feedforward and feedback control of arm dynamics in non-human primates. European Journal of Neuroscience 53, 1605–1620 (2021).

[36] Pei, F. et al. Neural latents benchmark ‘21: Evaluating latent variable models of neural population activity (2021). URL https://arxiv.org/abs/2109.04463.

[37] Keshtkaran, M. R. et al. A large-scale neural network training framework for generalized estimation of single-trial population dynamics. BioRxiv 2021–01 (2022).

[38] Perkins, S. M., Cunningham, J. P., Wang, Q. & Churchland, M. M. Simple decoding of behavior from a complicated neural manifold. bioRxiv 2023–04 (2023).

[39] Schneider, S., Lee, J. H. & Mathis, M. W. Learnable latent embeddings for joint behavioural and neural analysis. Nature 1–9 (2023).

[40] Hurwitz, C., Kudryashova, N., Onken, A. & Hennig, M. H. Building population models for large-scale neural recordings: Opportunities and pitfalls. Current opinion in neurobiology 70, 64–73 (2021).

[41] Churchland, M. M. et al. Neural population dynamics during reaching. Nature 487, 51–56 (2012).

