## Supplementary figures and tables for "A transient neural code for feedback-driven motor corrections during reaching"

**Table A1** Oscillating neurons in M1 vs PMd

| Monkey | Date | M1 neuron ids | PMd neuron ids | M1 count | PMd count |
|---|---|---|---|---|---|
| Chewie | 2016-09-15 | [15, 30, 31, 45] | [157] | 4 | 1 |
| Chewie | 2016-09-21 | [3, 27, 51] | [] | 3 | 1 |
| Chewie | 2016-10-05 | [30, 44] | [] | 2 | 0 |
| Chewie | 2016-10-07 | [1, 27, 23, 35] | [144, 157] | 4 | 2 |
| Chewie | total | | | 13 | 4 |
| Mihili | 2014-02-03 | [13] | [51] | 1 | 1 |
| Mihili | 2014-02-17 | [17] | [] | 1 | 0 |
| Mihili | 2014-02-18 | [5, 15] | [] | 2 | 0 |
| Mihili | 2014-03-07 | [2] | [] | 1 | 0 |
| Mihili | total | | | 5 | 1 |

### Appendix A Single neuron encoding of movement corrections

We used the Fourier transform and searched for a 4-5 Hz oscillations in neural firing rates of individual PMd and M1 neurons. We found a very small fraction of neurons with significant 4-5 Hz component in M1 (4±1%), and very few such neurons in PMd (0.2±0.5%, see Table A1). Most of the sessions did not contain any recorded oscillating PMd neurons. This suggests that M1 neurons have neural dynamic modes at a relevant timescale, and therefore the phase of oscillations in these individual M1 neurons might encode the phase of hand velocity oscillation.

Since the fraction of such neurons is small, this provides little evidence for single neurons capturing the observed oscillations. As we show in the paper based on the decoding analysis, the information about the phase and intensity of hand velocity oscillations is present in the M1 population code.

### Appendix B Hand velocity oscillations can be found in adaptation trials of all sessions

**Table B2** Datasets Information

| Dataset | | # trials | | | | | | # neurons | | |
|---|---|---|---|---|---|---|---|---|---|---|
| Monkey | Date | BL | AD | WO | Train | Test | # Time bins | All | M1 | PMd |
| Chewie | 2016-09-15 | 182 | 265 | 260 | 566 | 141 | 102 | 309 | 76 | 233 |
| Chewie | 2016-09-21 | 176 | 206 | 253 | 508 | 127 | 103 | 303 | 72 | 231 |
| Chewie | 2016-10-05 | 202 | 219 | 192 | 490 | 123 | 100 | 244 | 81 | 163 |
| Chewie | 2016-10-07 | 170 | 201 | 210 | 465 | 116 | 101 | 207 | 70 | 137 |
| Mihili | 2014-02-03 | 182 | 269 | 153 | 483 | 121 | 92 | 114 | 34 | 80 |
| Mihili | 2014-02-17 | 208 | 281 | 404 | 714 | 179 | 85 | 148 | 44 | 104 |
| Mihili | 2014-02-18 | 225 | 302 | 383 | 728 | 182 | 86 | 159 | 38 | 121 |
| Mihili | 2014-03-07 | 216 | 337 | 225 | 622 | 156 | 87 | 92 | 26 | 66 |

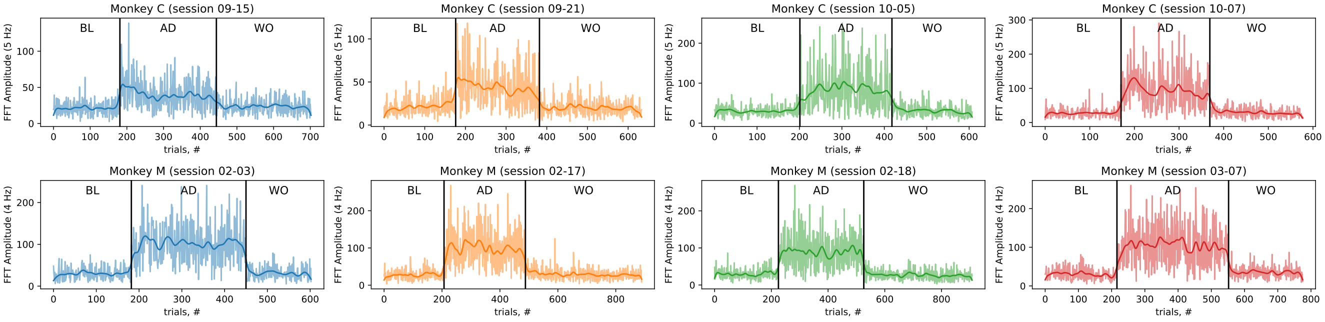


**Fig. B2** Fourier amplitude of hand velocity oscillation across trials and epochs, for both monkeys and all recording sessions.

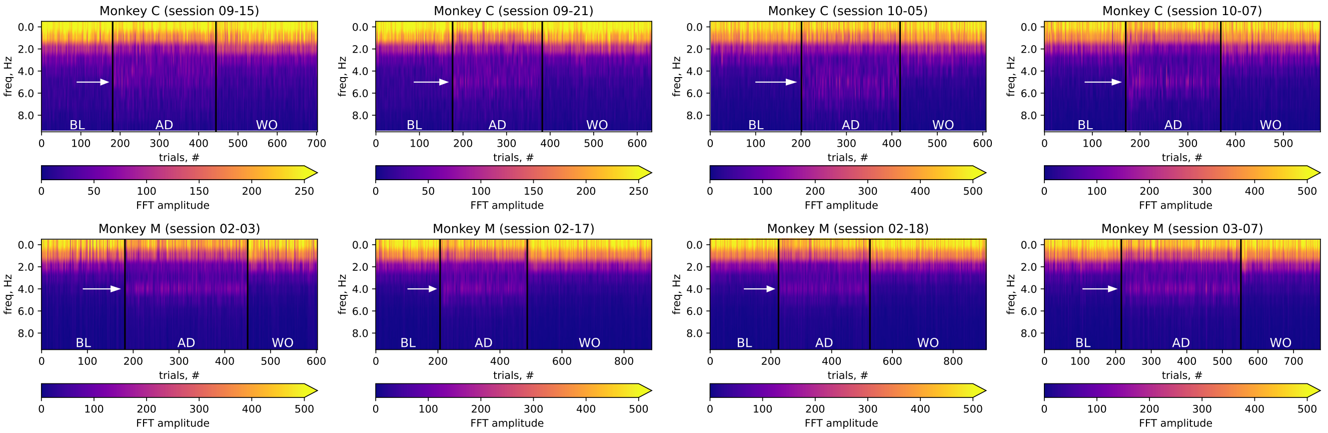


**Fig. B1** Fourier spectra of hand velocity across trials and epochs, for both monkeys and all recording sessions. The white arrow points at the oscillatory frequency (5 Hz for Monkey C, 4 Hz for Monkey M), which appears in adaptation trials.

| MC_Maze | Data resolution | 20ms bins | | | | 5ms bins | | | |
|---|---|---|---|---|---|---|---|---|---|
| 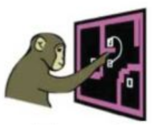 | Performance on NLB [2] | vel rank | vel $R^2$ | co-bps rank | co-bps | vel rank | vel $R^2$ | co-bps rank | co-bps |
| | **Our method (BAND)** | **#1** | **92.52%** | #3 | 0.3537 | **#1** | **93.62%** | #14 | 0.3215 |
| | CEBRA-behavior [4] | #2 | 91.06% | #9 | -1.3571 | – | | – | |
| | Churchland lab MINT [6] | #3 | 90.05% | #4 | 0.3295 | #2 | 91.21% | #12 | 0.3304 |
| | LFADS baseline [3] | #4 | 89.06% | **#2** | **0.3554** | #5 | 90.99% | **#10** | **0.3364** |
| **MC_RTT** | **Data resolution** | **20ms bins** | | | | **5ms bins** | | | |
| 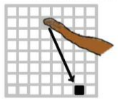 | **Performance on NLB [2]** | vel rank | vel $R^2$ | co-bps rank | co-bps | vel rank | vel $R^2$ | co-bps rank | co-bps |
| | **Our method (BAND)** | **#1** | **67.18%** | #3 | 0.1920 | **#1** | **67.05%** | #8 | 0.1846 |
| | Churchland lab MINT [6] | #2 | 65.47% | **#1** | **0.2008** | #2 | 65.59% | **#4** | **0.2014** |
| | LFADS baseline [3] | #3 | 61.05% | #2 | 0.1976 | #8 | 61.76% | #7 | 0.1868 |
| **Area2_Bump** | **Data resolution** | **20ms bins** | | | | **5ms bins** | | | |
| 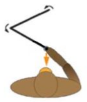 | **Performance on NLB [2]** | vel rank | vel $R^2$ | co-bps rank | co-bps | vel rank | vel $R^2$ | co-bps rank | co-bps |
| | **Our method (BAND)** | **#1** | **88.92%** | #4 | 0.2477 | **#1** | **89.50%** | #8 | 0.2686 |
| | Churchland lab MINT [6] | #2 | 88.03% | **#1** | **0.2718** | #2 | 88.77% | **#7** | **0.2735** |
| | LFADS baseline [3] | #4 | 85.65% | #3 | 0.2542 | #10 | 84.92% | #10 | 0.2569 |

Across all datasets, our method captured significant additional behavioral variability that corresponds to small neuronal variability.

#### C.1 Adapting BAND for NLB challenge

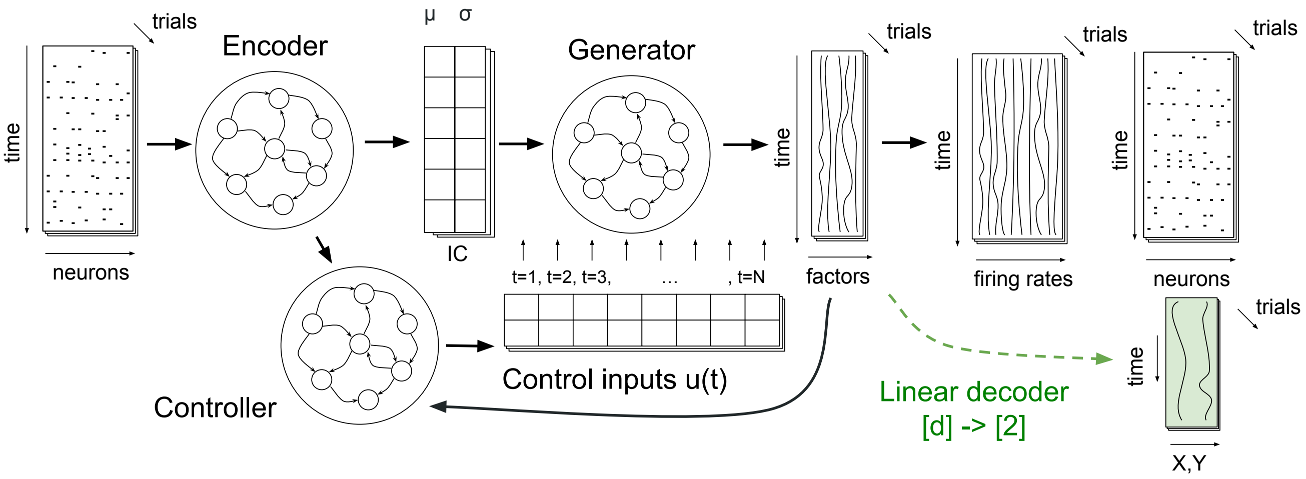


**Fig. C4** Acausal BAND archetecture with a fixed-lag behavior decoder for Neural Latents Benchmark challenge.

Here we utilized the exact same version of LFADS that demonstrated strong performance on the leaderboard. We used a simple linear behavior decoder, architecturally identical to the Ridge regression model used by NLB for evaluating behavioral $R^2$ (Fig. C4). While this version of the model was fine-tuned for the NLB challenge, its interpretability is limited, as well as its ability to capture behavioral feedback in neural activity due to the simplicity of the decoder.

We modified BAND decoder and objective in order to match the target metrics in NLB challenge. We used a standard linear decoder, instead of a sequence-to-sequence behavior decoder, to reproduce NLB decoding procedure. We also used the first two terms in BAND loss in (5) as a PBT objective for NLB challenge, with $\theta$ fixed to $10^{-4}$:

$$\mathcal{L}_{\text{PBT\_NLB}} = \mathcal{L}_x - 10^{-4} \cdot \text{MSE}(\mathbf{b}, \hat{\mathbf{b}}) \tag{C1}$$

Both PBT losses in (7) and (C1) are equivalent up to constant coefficients before neural and behavioral terms.

Note that there are multiple implementations of LFADS models in different frameworks: original in Tensorflow 1, Tensorflow 2, and PyTorch. We used Tensorflow 2 version for the official NLB challenge, while the rest of the paper uses PyTorch version.

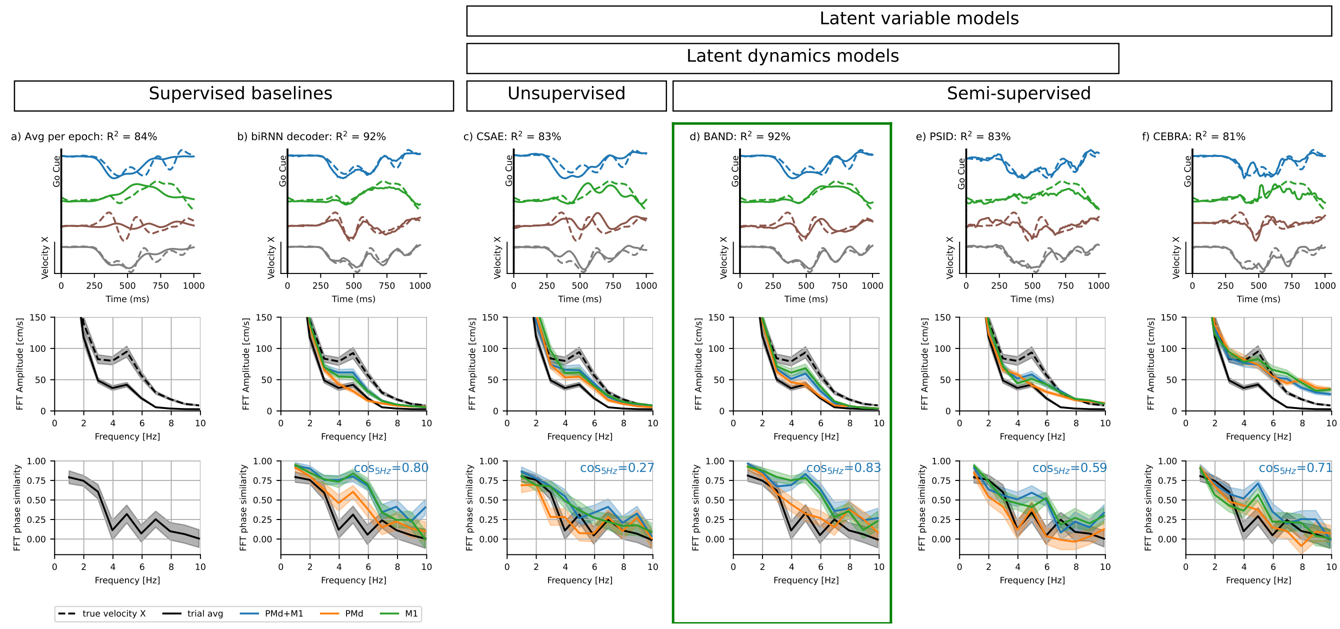


**Fig. C5** Model comparison shows that both nonlinear latent dynamics and behavior supervision are required for capturing hand velocity. Top row: hand velocity in example AD trials (solid line – model prediction from PMd+M1, dashed – ground truth); middle row: a Fourier spectrum of hand velocity predictions (from both or either brain areas), indicating whether higher amplitude of 5 Hz oscillations is correctly captured; bottom row: cosine similarity between Fourier modes of true velocity vs predicted velocity; cosine similarity at 5 Hz indicates whether the phase of oscillations is captured; **a)** An average velocity towards the reach target (average across all trials with the same reach target within AD epoch); **b)** Velocity predicted by a supervised bi-directional RNN decoder, demonstrating that hand velocity oscillations are decodable from neural activity; **c)** Velocity predicted using ridge regression from CSAE model with a 4-dimensional controller and 100 latent factors; **d)** Velocity predicted by BAND model with 100 factors and all hyperparameters matched to a CSAE model; **e)** Velocity predicted using a kNN decoder from CEBRA embedding; **f)** Velocity predicted by PSID model (a linear Kalman filter with neural and behavioral observations). Decoding here was performed on both PMd and M1 using all epochs, and results evaluated on adaptation epoch; predictions for all reach directions and individual brain areas can be found in Fig. H13)

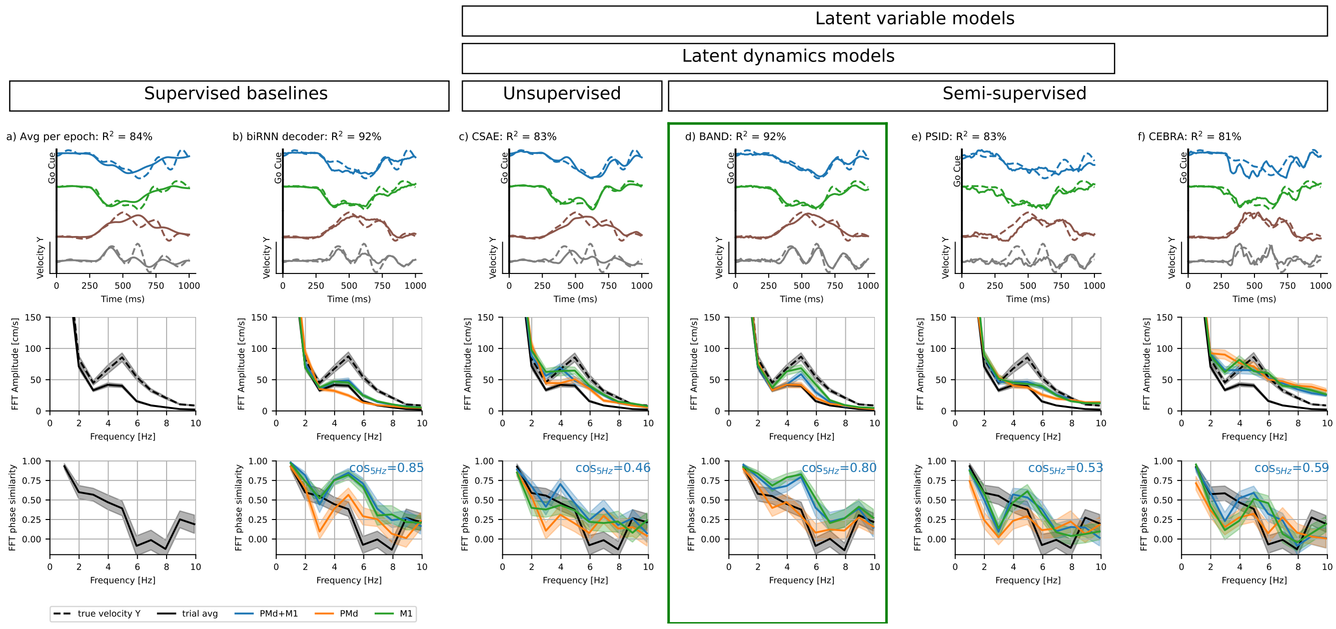


**Fig. C6** Same as Figure C5, but with velocity component Y.

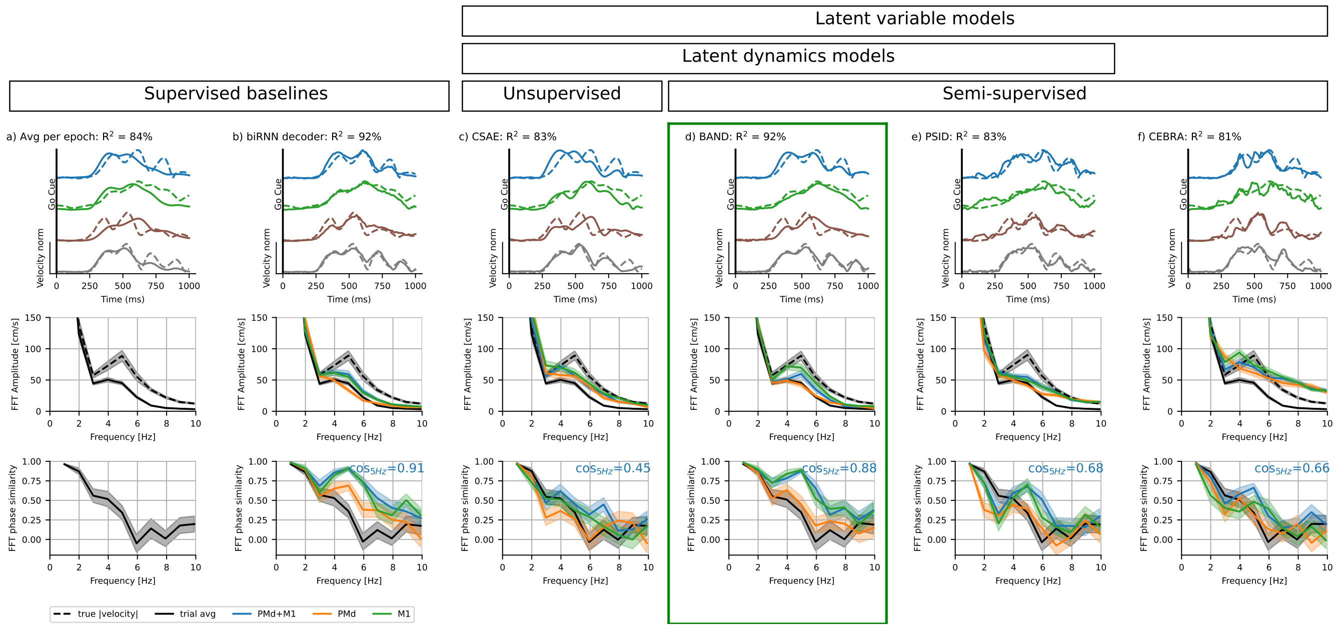


**Fig. C7** Same as Figure C5, but with the norm of hand velocity.

significant loss of neural reconstruction performance compared to CSAE (0.264 vs. 0.259 bits / second for the models shown here; standard deviation for cross-validated neural reconstruction is 0.004 bits / second). The fact that a CSAE model optimized for neural reconstruction quality discards these feedback-driven motor corrections suggests that they are encoded in low-variance patterns of neural activity.

### Appendix D Testing autonomous models (no controller trained)

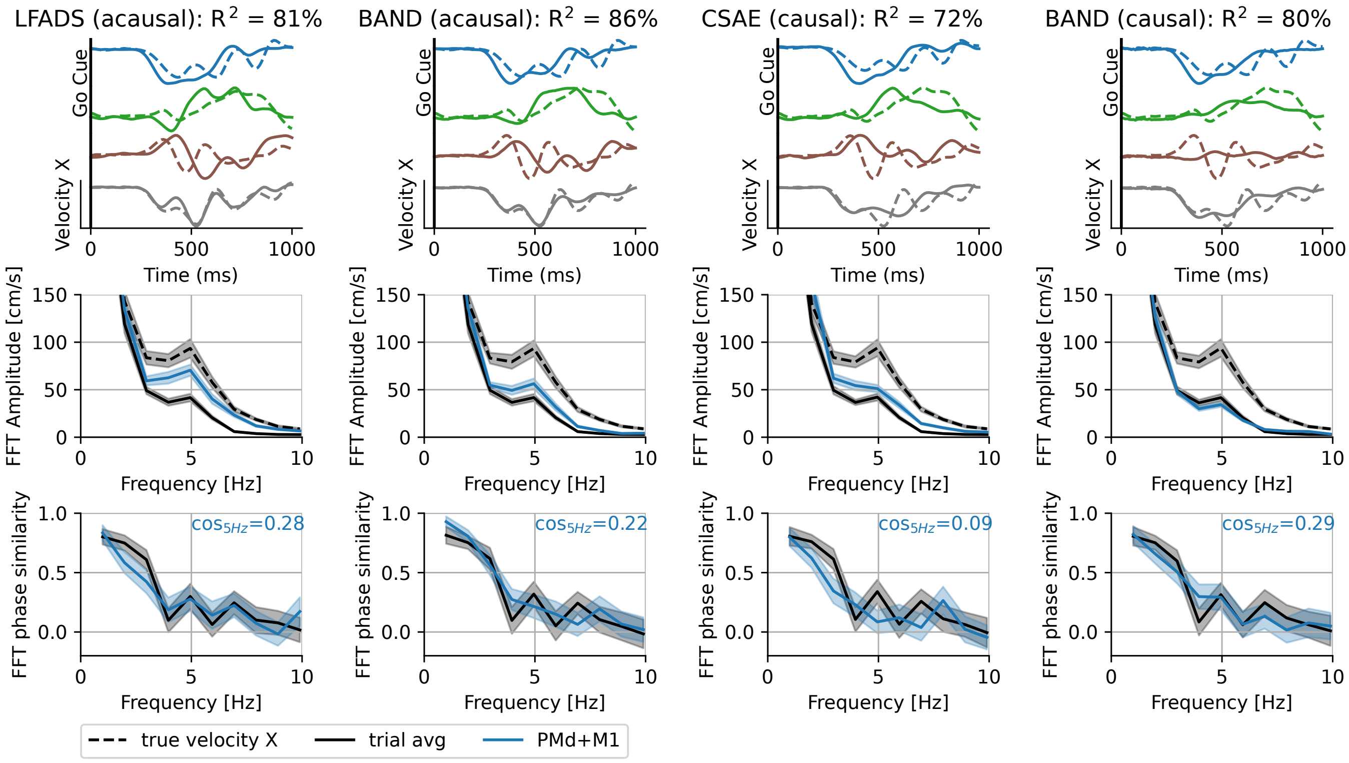


**Fig. D8** Model comparison for **autonomous** LFADS/BAND models, i.e. with controller removed from the model architecture.

Here we test whether autonomous models, i.e. without a controller, can learn latent factors predictive of hand velocity oscillations. This test demonstrates that the **causal** restrictions applied on our model prevent the model from compressing all the information about the trial into initial condition – a possible solution in original acausal LFADS, which lacks interpretability.

None of the models could predict the phase of oscillations correctly, even though acausal models had access to the future spiking data (bottom, Fig. D8). However, tuning all hyperparameters in the acausal models, as it is done in AutoLFADS [37], results in poorly regularized models with still create 5 Hz oscillations (middle row, Fig. D8), but fail to synchronize the phase. Adding causality prevents autonomous models from creating spurious oscillations.

### Appendix E Control inputs in BAND cause only a transient deviation from an autonomous trajectory

Inputs inferred via BAND, despite their transient nature, are not sparse in time; rather, they continually modulate the dynamics. By isolating individual inputs, we show that each input induces transient deviations from an autonomous trajectory without diverging from the long-term trajectory.

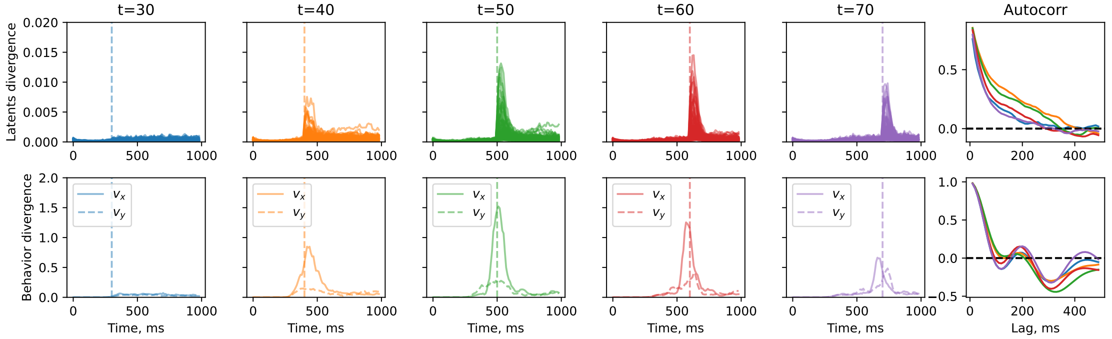


**Fig. E9** Deviation from the autonomous trajectory in the latent space and behavioral output space. Each line in the first five columns shows the trial-average residual along one of the dimensions. The last column shows autocorrelograms for the residuals evoked by perturbations.

By masking inputs in all but one time bin, we assessed how single-point perturbations affect latent and behavioral trajectories (Fig. E9). The resulting deviations (measured as a square distance) were transient, lasting less than 200 ms (see autocorrelogram half-width).

### Appendix F Control inputs in BAND correlate with movement speed

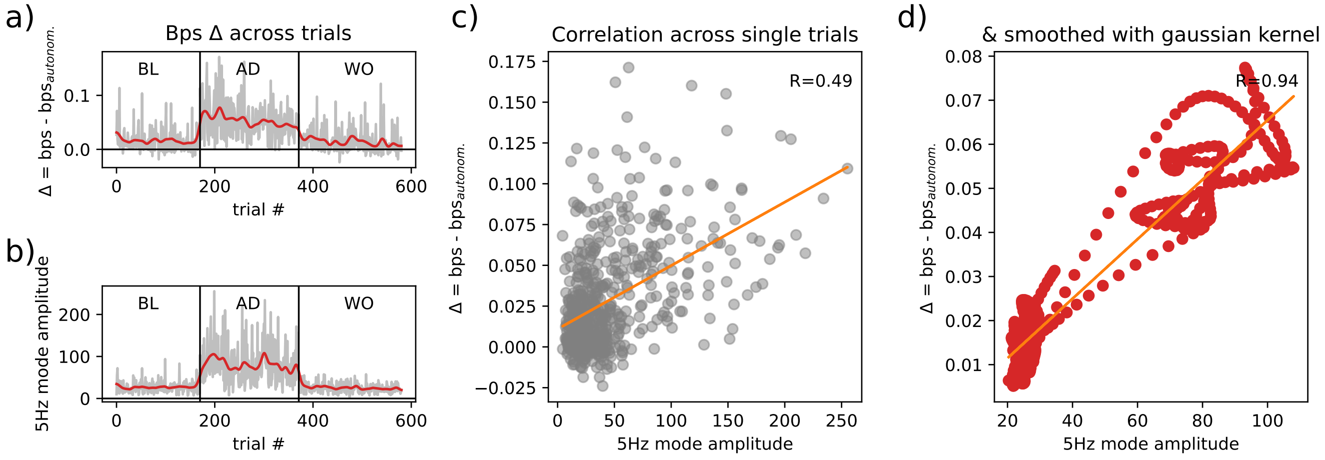


**Fig. F10** The controller adds oscillations to neural factors. a) BAND controller explains more neural variability (bits / second) in adaptation trials than in other epochs without perturbation; b) amplitude of the 5 Hz hand velocity oscillations (reproduces Fig. 2e, left); c) Correlation between single trial neural variability explained by controller (bps) and amplitude of the 5 Hz hand velocity oscillation mode (R=0.49); d) Correlation between across-trials smoothed trends (gaussian kernel s.d.=5 trials) of neural variability explained by controller (bps) and amplitude of the 5 Hz hand velocity oscillation mode (R=0.94);

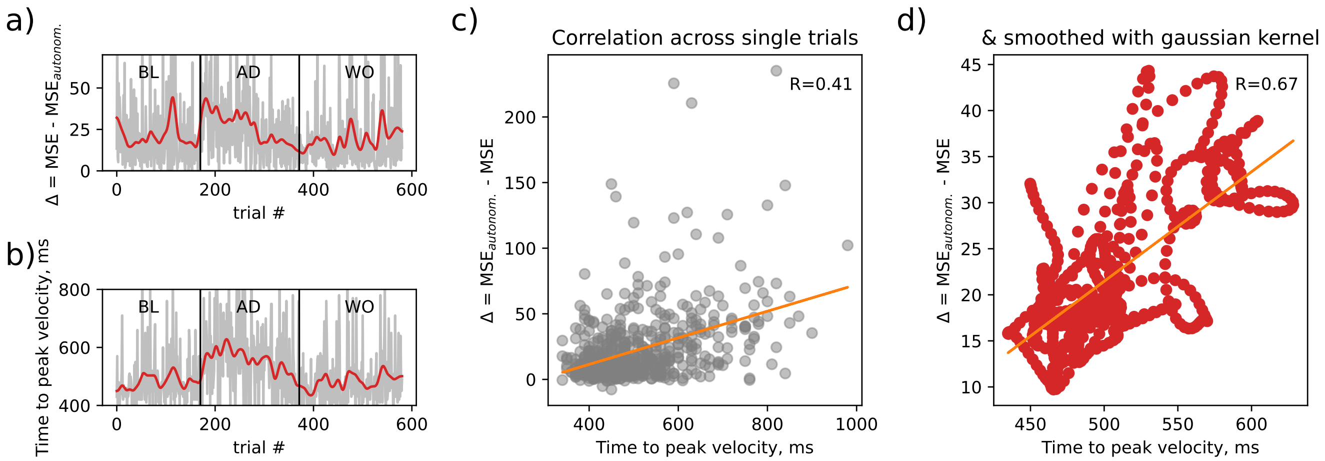


**Fig. F11** The controller contributes more to behavior prediction in the trials when reaching max velocity was delayed. a) Drop in mean-squared error due to controller ablation in BAND; b) Time from the start of the trial (movement onset - 250 ms) to reaching peak velocity; c) Correlation between single trial ΔMSE and time to reach peak velocity (R=0.41); d) Correlation between across-trials smoothed trends (gaussian kernel s.d.=5 trials) of ΔMSE and time to reach peak velocity (R=0.67);

### Appendix G Visualization of BAND latents

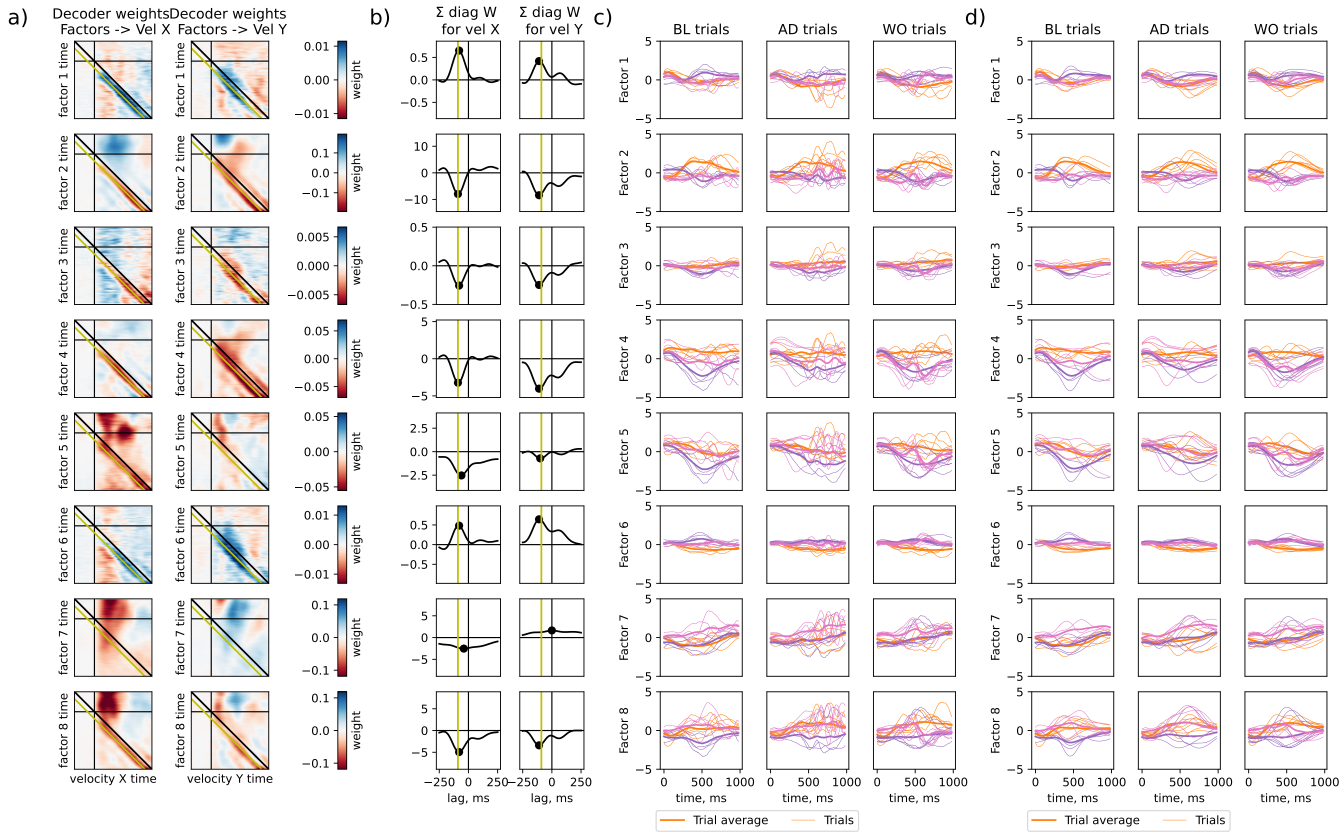


**Fig. G12** Behavior decoder weights and factors for a 8-factor BAND model with a causal controller.

### Appendix H Decoding performances ($R^2$ and cosine similarity) and trajectory divergence for different areas, methods, and sessions

**Table H3** Variance explained by an average hand velocity towards a given target ($R^2$, %)

| Monkey | Date | $R^2_{all}$ | $R^2_{BL}$ | $R^2_{AD}$ | $R^2_{WO}$ |
|---|---|---|---|---|---|
| Chewie | 2016-09-15 | 85.3 | 89.9 | 82.5 | 83.4 |
| Chewie | 2016-09-21 | 85.9 | 87.6 | 82.6 | 86.4 |
| Chewie | 2016-10-05 | 89.0 | 90.8 | 86.1 | 90.0 |
| Chewie | 2016-10-07 | 86.8 | 84.6 | 83.8 | 90.7 |
| Mihili | 2014-02-03 | 79.9 | 82.8 | 81.1 | 74.4 |
| Mihili | 2014-02-17 | 81.8 | 83.9 | 81.2 | 80.6 |
| Mihili | 2014-02-18 | 86.9 | 88.8 | 88.0 | 84.9 |
| Mihili | 2014-03-07 | 84.1 | 85.9 | 82.6 | 83.9 |

| Monkey | Date | both_$R^2_{all}$ | both_$R^2_{BL}$ | both_$R^2_{AD}$ | both_$R^2_{WO}$ | PMd_$R^2_{all}$ | PMd_$R^2_{BL}$ | PMd_$R^2_{AD}$ | PMd_$R^2_{WO}$ | M1_$R^2_{all}$ | M1_$R^2_{BL}$ | M1_$R^2_{AD}$ | M1_$R^2_{WO}$ |
|---|---|---|---|---|---|---|---|---|---|---|---|---|---|
| Chewie | 2016-09-15 | 93.2 | 94.6 | 90.8 | 93.9 | 91.1 | 92.8 | 88.3 | 91.6 | 91.1 | 93.1 | 87.8 | 91.8 |
| Chewie | 2016-09-21 | 92.6 | 93.9 | 88.7 | 93.9 | 90.2 | 91.7 | 85.0 | 91.9 | 90.2 | 91.4 | 86.3 | 91.5 |
| Chewie | 2016-10-05 | 94.9 | 96.2 | 92.9 | 95.5 | 91.2 | 93.0 | 87.7 | 92.7 | 93.6 | 95.3 | 91.0 | 94.6 |
| Chewie | 2016-10-07 | 94.2 | 94.6 | 92.0 | 95.3 | 90.0 | 92.0 | 84.6 | 92.3 | 93.2 | 93.0 | 90.8 | 94.9 |
| Mihili | 2014-02-03 | 85.6 | 85.9 | 85.1 | 84.8 | 73.6 | 73.9 | 75.2 | 70.2 | 82.1 | 82.1 | 81.0 | 81.7 |
| Mihili | 2014-02-17 | 91.8 | 93.5 | 88.7 | 92.2 | 84.4 | 87.8 | 81.4 | 83.8 | 85.6 | 86.4 | 83.3 | 86.0 |
| Mihili | 2014-02-18 | 91.8 | 92.2 | 89.4 | 92.8 | 87.5 | 90.3 | 84.7 | 87.3 | 86.8 | 88.3 | 82.1 | 88.4 |
| Mihili | 2014-03-07 | 87.8 | 89.0 | 85.3 | 88.7 | 82.5 | 84.2 | 80.2 | 82.3 | 83.7 | 85.6 | 81.1 | 83.8 |

**Table H4** Variance explained by a biRNN decoder given neural recordings from M1, PMd, or both areas together ($R^2$, %). Red indicates cases in which biRNN decoding captured less variance than a model that only predicts reach direction (i.e. $R^2$ is below that in Table H3).

| Monkey | Date | both_$R^2_{all}$ | both_$R^2_{BL}$ | both_$R^2_{AD}$ | both_$R^2_{WO}$ | PMd_$R^2_{all}$ | PMd_$R^2_{BL}$ | PMd_$R^2_{AD}$ | PMd_$R^2_{WO}$ | M1_$R^2_{all}$ | M1_$R^2_{BL}$ | M1_$R^2_{AD}$ | M1_$R^2_{WO}$ |
|---|---|---|---|---|---|---|---|---|---|---|---|---|---|
| Chewie | 2016-09-15 | 84.3 | 84.4 | 80.9 | 86.3 | 79.0 | 80.6 | 75.0 | 80.3 | 66.7 | 71.1 | 49.8 | 75.0 |
| Chewie | 2016-09-21 | 79.3 | 80.5 | 73.4 | 81.4 | 75.1 | 77.3 | 70.0 | 76.0 | 67.6 | 69.1 | 59.5 | 70.8 |
| Chewie | 2016-10-05 | 84.1 | 85.2 | 81.1 | 85.8 | 75.1 | 76.0 | 71.7 | 77.6 | 76.7 | 80.2 | 72.3 | 77.2 |
| Chewie | 2016-10-07 | 82.8 | 80.7 | 80.6 | 85.9 | 72.6 | 74.3 | 67.6 | 74.6 | 71.7 | 66.8 | 71.0 | 75.4 |
| Mihili | 2014-02-03 | 58.7 | 57.9 | 58.2 | 56.6 | 30.0 | 33.5 | 19.4 | 31.5 | 43.8 | 49.7 | 38.4 | 36.9 |
| Mihili | 2014-02-17 | 71.8 | 76.3 | 67.4 | 70.9 | 52.4 | 59.3 | 46.3 | 50.6 | 54.5 | 60.5 | 45.6 | 55.1 |
| Mihili | 2014-02-18 | 73.4 | 72.6 | 72.6 | 73.5 | 58.6 | 58.2 | 59.2 | 57.2 | 55.8 | 56.8 | 52.5 | 56.3 |
| Mihili | 2014-03-07 | 68.3 | 67.9 | 69.0 | 66.0 | 42.3 | 37.9 | 37.2 | 48.7 | 49.5 | 50.2 | 50.8 | 44.7 |

**Table H5** Variance explained by a CEBRA + kNN decoder given neural recordings from M1, PMd, or both areas together ($R^2$, %). One green value indicates cases in which CEBRA explained more variance than the average hand trajectory (i.e. Table. H3).

We applied Fourier decomposition to hand velocity predictions from biRNN decoder (Fig. H13), LFADS (Fig. H14) and BAND (Fig. H16).

| Monkey | Date | both_$R^2_{all}$ | both_$R^2_{BL}$ | both_$R^2_{AD}$ | both_$R^2_{WO}$ | PMd_$R^2_{all}$ | PMd_$R^2_{BL}$ | PMd_$R^2_{AD}$ | PMd_$R^2_{WO}$ | M1_$R^2_{all}$ | M1_$R^2_{BL}$ | M1_$R^2_{AD}$ | M1_$R^2_{WO}$ |
|---|---|---|---|---|---|---|---|---|---|---|---|---|---|
| Chewie | 2016-09-15 | 82.9 | 85.2 | 78.1 | 84.4 | 77.8 | 80.7 | 74.1 | 78.2 | 78.0 | 81.7 | 66.4 | 83.1 |
| Chewie | 2016-09-21 | 80.9 | 84.5 | 74.6 | 81.7 | 74.9 | 77.4 | 67.9 | 76.8 | 77.9 | 83.3 | 69.6 | 78.6 |
| Chewie | 2016-10-05 | 82.5 | 84.4 | 81.4 | 81.6 | 71.9 | 72.4 | 71.0 | 72.2 | 83.3 | 86.2 | 80.5 | 83.2 |
| Chewie | 2016-10-07 | 83.9 | 84.8 | 82.6 | 83.7 | 72.9 | 72.7 | 68.6 | 76.0 | 80.8 | 83.4 | 79.2 | 79.1 |
| Mihili | 2014-02-03 | 74.0 | 75.6 | 71.9 | 72.3 | 52.0 | 51.9 | 48.5 | 51.9 | 70.9 | 76.0 | 67.2 | 67.4 |
| Mihili | 2014-02-17 | 81.7 | 84.2 | 76.4 | 82.8 | 70.0 | 70.4 | 66.3 | 70.8 | 76.5 | 80.2 | 68.9 | 77.9 |
| Mihili | 2014-02-18 | 83.9 | 83.5 | 80.9 | 85.8 | 75.0 | 73.7 | 75.5 | 75.3 | 78.1 | 80.9 | 71.3 | 80.1 |
| Mihili | 2014-03-07 | 78.6 | 78.8 | 76.0 | 79.7 | 66.1 | 65.4 | 64.5 | 67.0 | 71.2 | 71.6 | 68.1 | 72.5 |

**Table H6** Variance explained by a PSID decoder given neural recordings from M1, PMd, or both areas together ($R^2$, %). Green values indicate cases in which PSID explained more variance than the average hand trajectory (i.e. Table. H3). Note, that PSID never explained uninstructed variance in perturbed trials.

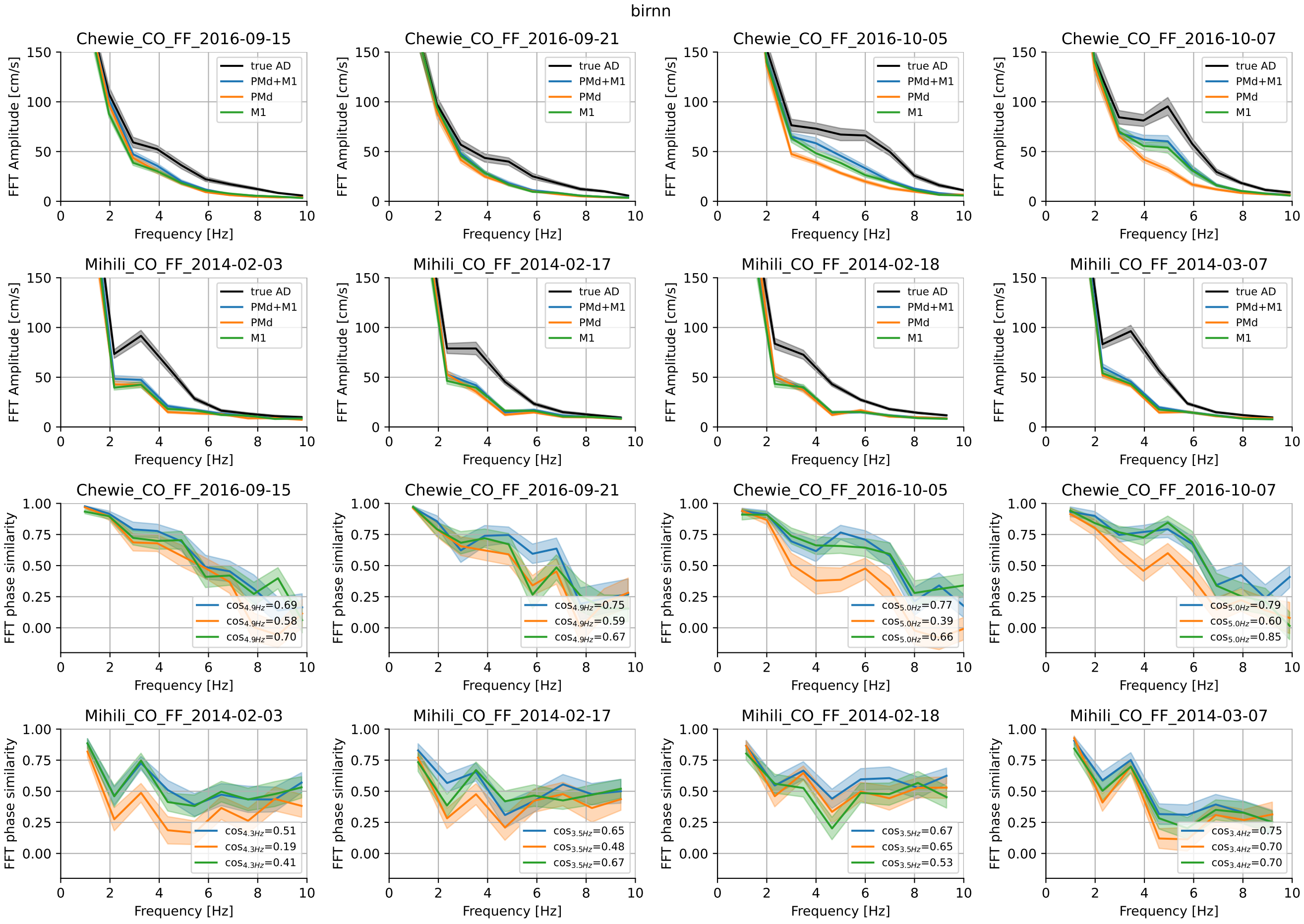


**Fig. H13** Fourier spectrum for velocity predictions based on biRNN decoder. In the sessions where hand velocity oscillations are decodable, they are decodable from M1 and not PMd. Note, that the number of M1 neurons recorded in Monkey M is considerably lower then in Monkey C. Brain areas used in the decoding are color-coded (blue: PMd+M1, orange: PMd, green: M1).

### References


[1] Churchland, M. M., Afshar, A. & Shenoy, K. V. A central source of movement variability. *Neuron* **52**, 1085–1096 (2006).

[2] Vyas, S., Golub, M. D., Sussillo, D. & Shenoy, K. V. Computation through neural population dynamics. *Annual review of neuroscience* **43**, 249–275 (2020).

[3] Kao, T.-C., Sadabadi, M. S. & Hennequin, G. Optimal anticipatory control as a theory of motor preparation: A thalamo-cortical circuit model. *Neuron* **109**, 1567–1581 (2021).

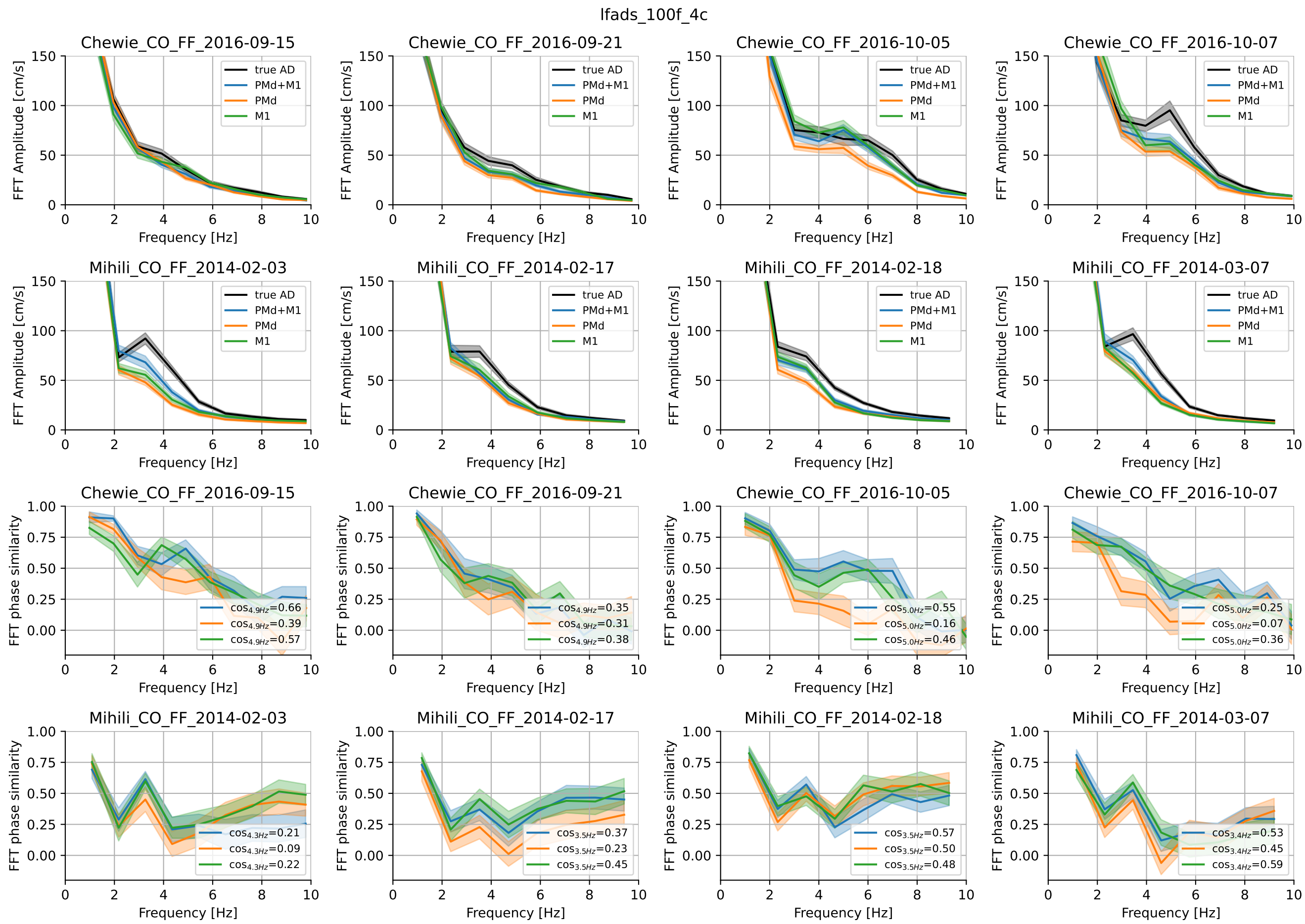


**Fig. H14** Fourier spectrum for velocity predictions based on CSAE models with 100 factors.


[4] Schimel, M., Kao, T.-C., Jensen, K. T. & Hennequin, G. ilqr-vae: control-based learning of input-driven dynamics with applications to neural data. *bioRxiv* 2021–10 (2021).

[5] Pandarinath, C. *et al.* Inferring single-trial neural population dynamics using sequential auto-encoders. *Nature methods* **15**, 805–815 (2018).

[6] Perich, M. G., Gallego, J. A. & Miller, L. E. A neural population mechanism for rapid learning. *Neuron* **100**, 964–976 (2018).

[7] Gandolfo, F., Li, C.-S., Benda, B., Schioppa, C. P. & Bizzi, E. Cortical correlates of learning in monkeys adapting to a new dynamical environment. *Proceedings of the National Academy of Sciences* **97**, 2259–2263 (2000).

[8] Li, C.-S. R., Padoa-Schioppa, C. & Bizzi, E. Neuronal correlates of motor performance and motor learning in the primary motor cortex of monkeys adapting to an external force field. *Neuron* **30**, 593–607 (2001).

[9] Arce, F., Novick, I., Mandelblat-Cerf, Y. & Vaadia, E. Neuronal correlates of memory formation in motor cortex after adaptation to force field. *Journal of Neuroscience* **30**, 9189–9198 (2010).

[10] Sun, X. *et al.* Cortical preparatory activity indexes learned motor memories. *Nature* **602**, 274–279 (2022).

[11] Dooley, J. C. & Blumberg, M. S. Developmental'awakening'of primary motor cortex to the sensory consequences of movement. *elife* **7**, e41841 (2018).

[12] Hurwitz, C. *et al.* Targeted neural dynamical modeling. *NeurIPS* **34**, 29379–29392 (2021).

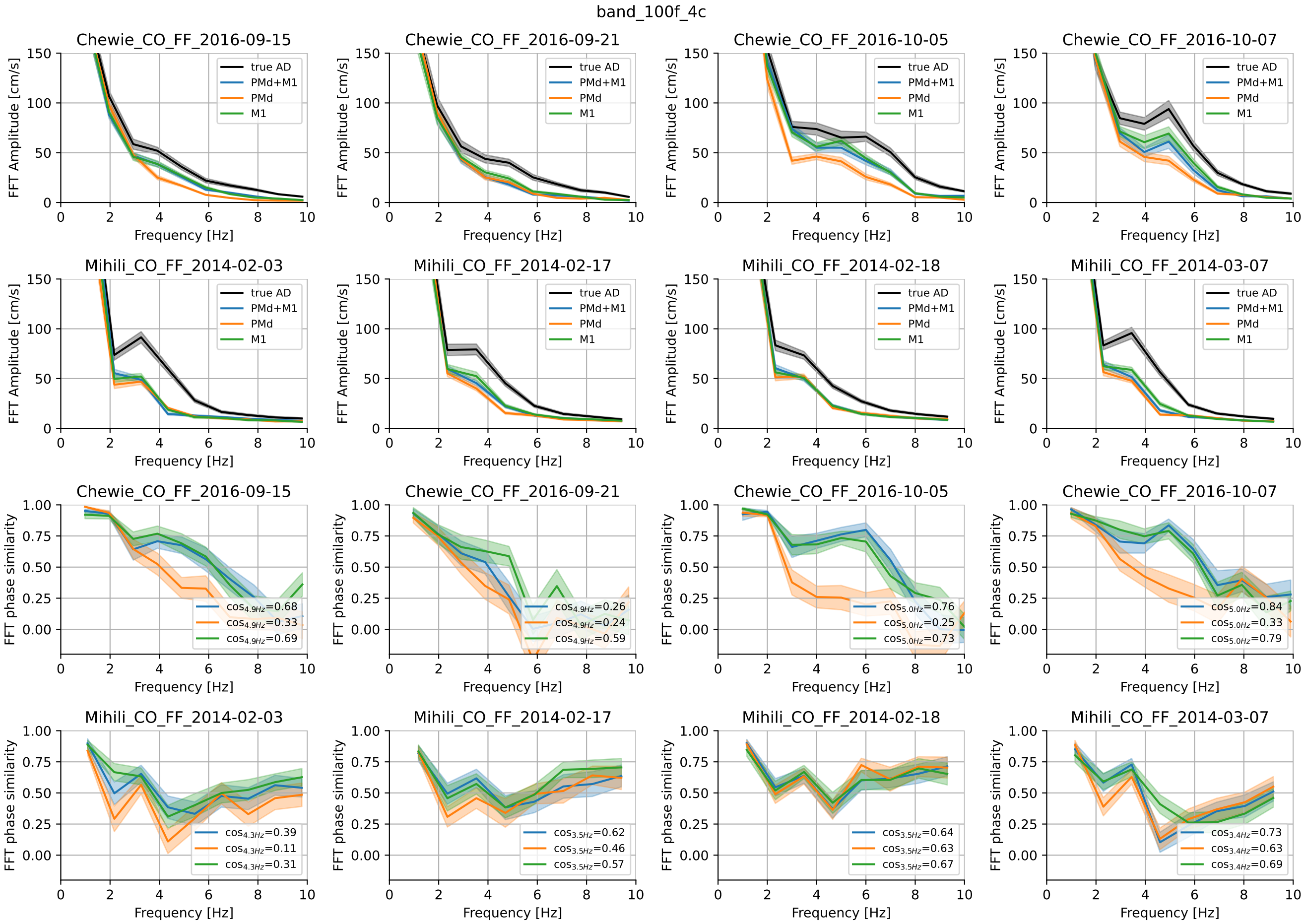


**Fig. H15** Fourier spectrum for velocity predictions based on BAND models with 100 factors.


[13] Georgopoulos, A., Kalaska, J., Caminiti, R. & Massey, J. Interruption of motor cortical discharge subserving aimed arm movements. *Experimental Brain Research* **49**, 327–340 (1983).

[14] Pruszynski, J. A. *et al.* Primary motor cortex underlies multi-joint integration for fast feedback control. *Nature* **478**, 387–390 (2011).

[15] Scott, S. H. Optimal feedback control and the neural basis of volitional motor control. *Nature Reviews Neuroscience* **5**, 532–545 (2004).

[16] Versteeg, C. & Miller, L. Dynamical feedback control: motor cortex as an optimal feedback controller based on neural dynamics (2022).

[17] Gurnani, H., Liu, W. & Brunton, B. W. Feedback control of recurrent dynamics constrains learning timescales during motor adaptation. *bioRxiv* 2024–05 (2024).

[18] Michaels, J. A. *et al.* Sensory expectations shape neural population dynamics in motor circuits. *Nature* 1–10 (2025).

[19] Perich, M. G., Narain, D. & Gallego, J. A. A neural manifold view of the brain. *Nature Neuroscience* 1–16 (2025).

[20] Sani, O. G., Abbaspourazad, H., Wong, Y. T., Pesaran, B. & Shanechi, M. M. Modeling behaviorally relevant neural dynamics enabled by preferential subspace identification. *Nature Neuroscience* **24**, 140–149 (2021).

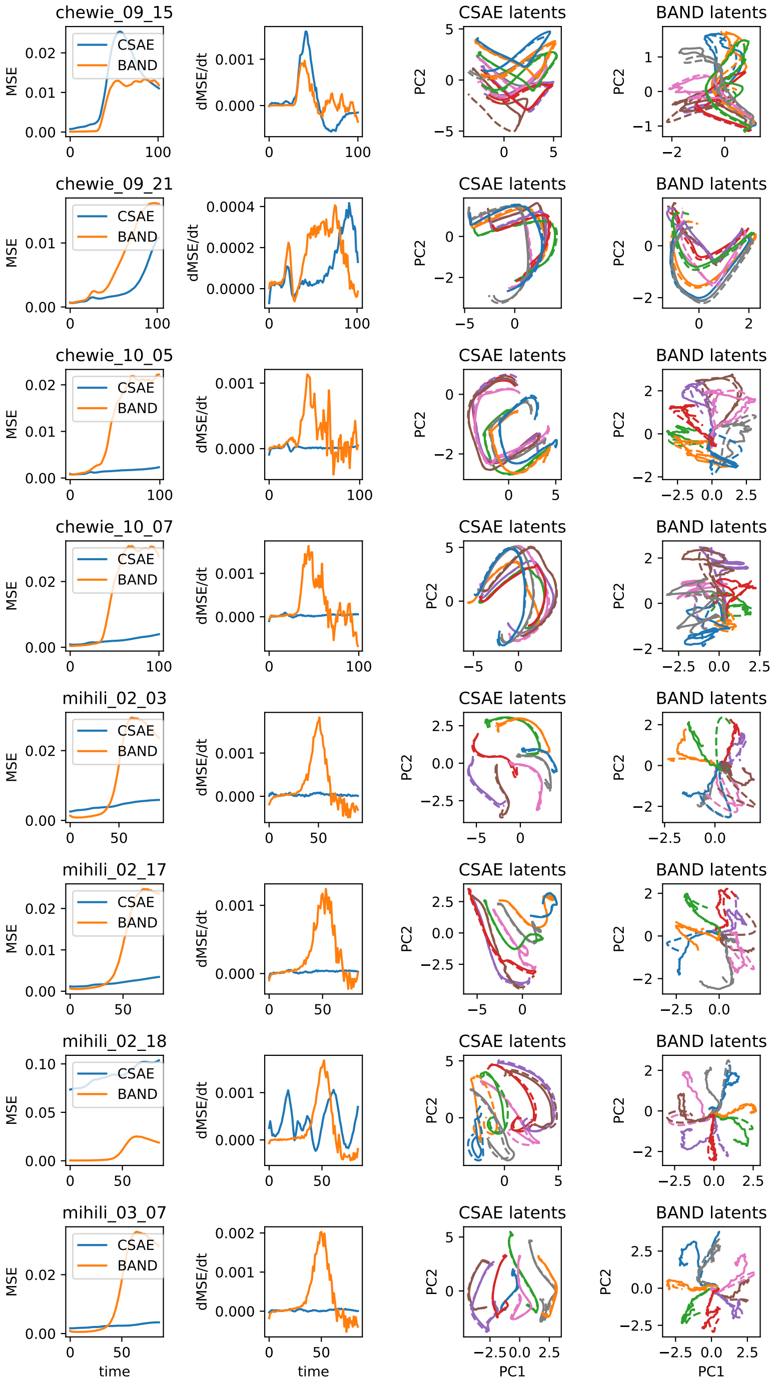


**Fig. H16** Latent trajectory divergence for controlled vs. autonomous latent trajectories in CSAE / BAND across animals and sessions. All neurons (M1 and PMd) were used in training these models, and all datasets were aligned to go cue. BAND shows consistent transient trajectory divergence during movement (orange traces), which then stabilizes (dMSE/dt=0) or even begins to converge (dMSE/dt¡0). CSAE demonstrates a variety of behaviors (blue traces), with most models slightly diverging from the autonomous trajectories towards the end.
